# BASCULE: Bayesian inference and clustering of mutational signatures leveraging biological priors

**DOI:** 10.1101/2024.09.16.613266

**Authors:** Elena Buscaroli, Azad Sadr, Riccardo Bergamin, Salvatore Milite, Edith Natalia Villegas Garcia, Arianna Tasciotti, Alessio Ansuini, Daniele Ramazzotti, Nicola Calonaci, Giulio Caravagna

**Affiliations:** Department of Mathematics, Informatics and Geosciences, University of Trieste, Trieste, Italy; Centre for Computational Biology, Human Technopole, Milan, Italy; Area Science Park, Trieste, Italy; University of Milan-Bicocca, Italy

## Abstract

Mutational signatures provide key insights into cancer mutational processes, but the availability of signature catalogues generated by different groups using distinct methodologies underscores a need for standardisation. We introduce a Bayesian framework that offers a systematic approach to expanding existing signature catalogues for any type of mutational signature, while grouping patients based on shared signature patterns. We demonstrate that this approach can identify both known and novel molecular subtypes across nearly 8,000 samples spanning six cancer types, and show that stratifications derived from signature yield prognostic groups, further enhancing the translational potential of mutational signatures.

## Background

Clonal dynamics in tumours [1–3] are primarily determined by somatic mutations that accumulate during cell division [4–6]. These mutations can arise in various forms. Single Base Substitutions (SBS) and insertions/deletions (indels) are often caused by random DNA replication errors or impaired mismatch repair mechanisms [7–9], while more complex structural variants, such as copy number alterations (CNAs), result from chromosome segregation errors, breakage, and rejoining [10, 11]. Both endogenous and exogenous mutational processes influence the rate of these mutational events, and substantial efforts are underway to decipher the mechanisms underlying them.

One of the most notable results is the introduction, in 2013, of the concept of SBS signatures, an idea later extended to dinucleotide-base substitutions (DBS), small indels (ID) and eventually copy number (CN) signatures [12–15]. In an actual tumour, multiple mutational processes co-exist, depending on the molecular features, the patient’s genetic background, and environmental factors, including treatment [16]. Understanding these mutational processes has significant translational potential, as it can inform treatment strategies and improve patient outcomes. For instance, mutational signatures associated with impaired mismatch repair can guide the use of immune checkpoint inhibitors in patients with microsatellite instability-high tumours [17]. Similarly, mutational profiles linked to homologous recombination deficiency have been used to identify patients who may benefit from PARP inhibitors in breast and ovarian cancers [18]. These examples underscore how insights into mutational mechanisms can directly inform therapeutic decisions, advancing precision medicine in oncology [19, 20].

Computational tools are increasingly used to detect mutational signatures from mutation count data derived from whole-exome or whole-genome sequencing (WGS). These tools generally fall into two categories [21]: those that apply established signal deconvolution algorithms to extract mutational signatures and estimate their prevalence across samples, a process known as de novo signature discovery [22–27], and those that focus on decomposing mutation data using predefined reference signature catalogues, bypassing the need for de novo discovery [22, 23, 27, 28]. These methodologies have been instrumental in assembling mutational process catalogues in cancer [21], including the development of COSMIC (Catalogue of Somatic Mutations in Cancer), which currently lists 144 mutational signatures — 94 SBS, 11 DBS, 18 ID, and 21 CN signatures [29]. A subset of these signatures has been experimentally linked to specific endogenous and exogenous processes, shedding light on their underlying mechanisms [30]. For example, distinct SBS and DBS signatures have been associated with exposures such as tobacco smoking, UV light, platinum-based therapies, and asbestos, showing differences in strand bias, GC context, and flanking bases [20]. Similarly, defective homologous recombination in DNA repair and AID/APOBEC-mediated mutagenesis have given rise to SBS and ID signatures, while chromothripsis and genome duplication events have produced CN signatures. A recent analysis by [23] used primary and metastatic WGS samples to expand COSMIC with 40 additional SBS and 18 DBS signatures [23].

In this rapidly evolving field, the challenge lies in harmonising multiple mutational catalogues produced by different research groups and computational tools, a process often relying on similarity metrics such as cosine similarity. However, this task requires extensive human oversight and careful parameter optimization. Ideally, more advanced tools would leverage existing knowledge to streamline this process. A unified catalogue encompassing various signature types could aid in translating mutational signatures into a stratification metric, enabling the identification of molecular subtypes — groups of patients with similar mutational profiles. Unfortunately, no current tool is capable of performing de novo signature discovery while integrating known signature catalogues, nor can any tool stratify patients based on signature exposure.

We introduce BASCULE (Fig. 1A), a Bayesian framework that detects mutational signatures – of any kind – by leveraging an existing catalogue, ensuring that newly identified signatures are statistically distinct from known ones (Fig. 1B). Additionally, BASCULE groups patients based on similar mutational profiles by analysing exposure across various signature types. Both algorithms are implemented using stochastic variational inference, built in the Pyro probabilistic programming language [31]. This design enables BASCULE to run natively on GPUs, allowing for seamless scaling to handle large datasets efficiently. Moreover, we release BASCULE as a Nextflow [32] module that can be embedded in any pipeline via a Singularity image, enabling reproducible signatures analysis across platforms. After validating the tool through simulations, we applied it to SBS and DBS signatures from ∼7,000 samples published by [23] and SBS signatures from ∼1000 ICGC (International Cancer Genome Consortium) cohort samples. BASCULE successfully identified well-established molecular subtypes in breast, lung, and colorectal cancers, while also generating new hypotheses about previously unreported co-occurrences of SBS and DBS signatures in specific patient groups. Additionally we conducted survival analysis on skin, pancreatic and esophageal cancers, revealing distinct, well differentiated clusters in skin and pancreatic cancer types. Building on prior research, BASCULE offers a powerful approach to enhance our understanding of mutational processes in cancer and uncover novel molecular insights.

**Fig. 1.**
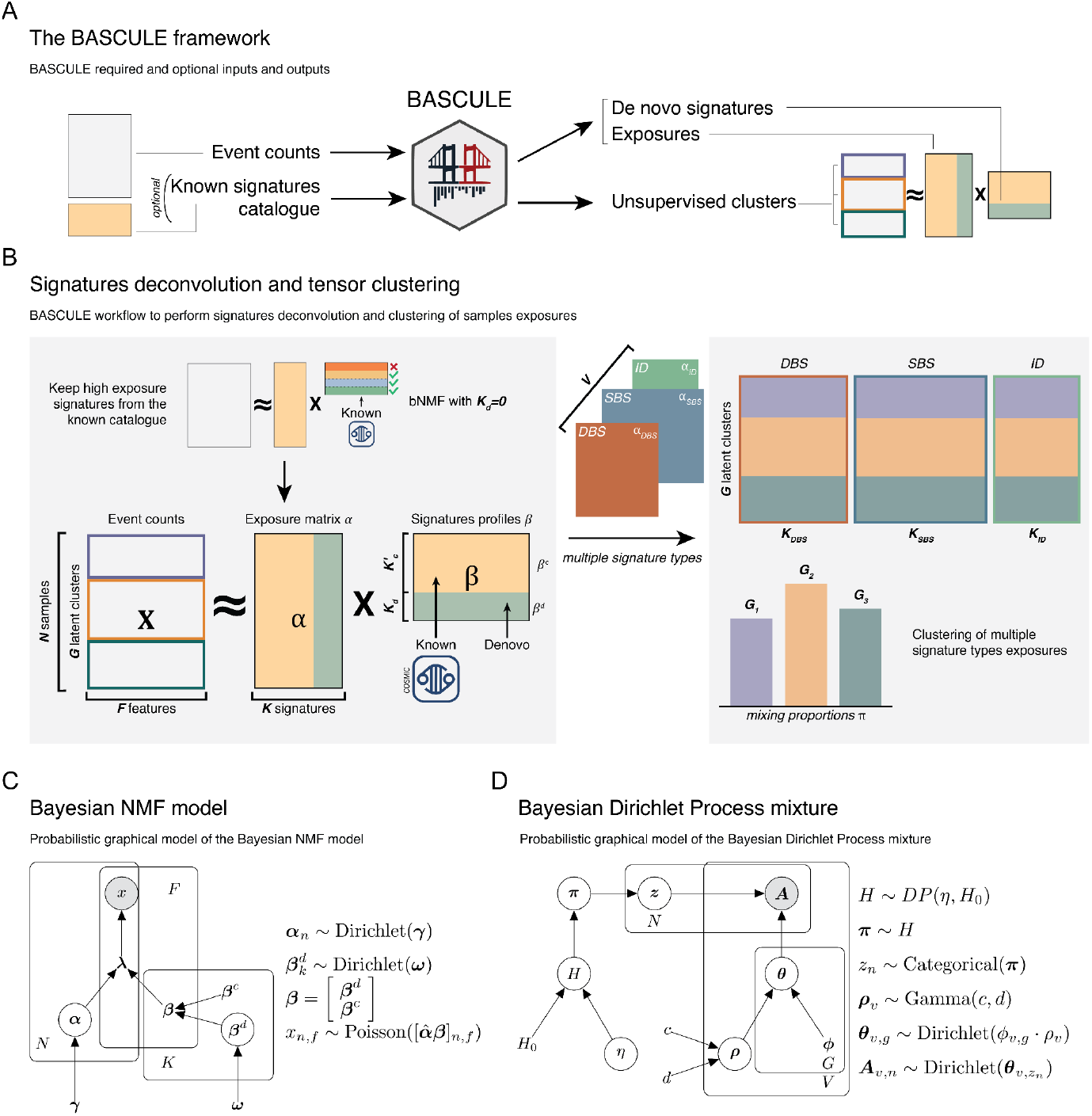
A. BASCULE decomposes an event counts matrix into signatures and their exposures while leveraging prior information; then, it clusters samples according to their exposures. B. BASCULE performs signatures deconvolution with a two-step inference process. First, it identifies relevant signatures from a catalogue of known signatures via Bayesian non-negative matrix factorisation. Then, it performs de novo signatures discovery, augmenting the catalogue signatures. BASCULE can deconvolute any signature type (SBS, DBS, IDS, CNS, etc.) and use their exposures to jointly cluster the samples. C. The Bayesian non-negative matrix factorisation model: the observed mutation counts matrix ***X*** follows a Poisson distribution, whose rate is defined by the product of the latent exposures *β* and signatures *β* matrices (Methods). D. BASCULE clusters via the Bayesian Dirichlet Process mixture model. The tensor of exposure matrices is distributed as a Dirichlet, along with its latent group-specific concentration *θ*. Samples are assigned to the cluster, maximising the posterior distribution (Methods).

## Results

### Bayesian inference of mutational signatures using a catalogue

BASCULE has a probabilistic model (Fig. 1C) that extracts a set *β* of signatures from data of *N* patients, along with signature exposures *α*. In what follows, we refer broadly to mutational signatures as anything that could be linked to somatic substitutions, regardless these are single-base substitutions or more complex signatures such as those derived from aneuploidy. The main novelty of BASCULE is that *β* is a combination of pre-specified catalogued signatures *β*^*c*^, and de novo signatures *β*^*d*^; signatures *β*^*d*^ in differ, probabilistically, from *β*^*c*^. The model is general and works with any signature type (SBS, DBS, ID or CN). To extract signatures, we use a Bayesian non-negative matrix factorization (bNMF) approach that factorises ***X***, a matrix of non-negative values for the observed counts detected in patients, as

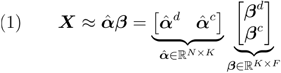

such that *K* =*K*^*d*^ + *K*^*c*^ The matrix 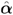 represents the number of mutations attributed to each mutational signature within a sample. Its normalised counterpart, *α*, expresses these contributions as proportions. Thus, 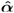 is obtained by multiplying the normalised exposures *α* by the total number of mutations in each sample, as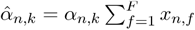, where 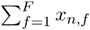 denotes the total mutation count for sample *n*. The matrices of *α* and *β* report multinomial random variables, that is

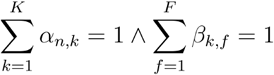

The model likelihood is Poisson

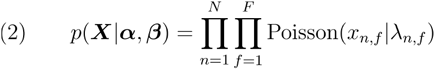

with rate defined by the reconstructed count matrix 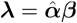.

In this model, the columns *F* are features that depend on the type of signature (e.g., for SBS and DBS these are 96 and 78 strand-agnostic substitution contexts, respectively), and *X* is decomposed into two pairs of exposure and signature matrices, one with *K*^*c*^ signatures from the catalogue, and one with *K*^*d*^ de novo signatures. We note that in the learning process, the signatures from the catalogue *β* ^*c*^ are not learnt from the data, i.e., they are held fixed based on the catalogue. Instead, the exposure vectors are learnt for both catalogue and de novo signatures. The overall bNMF joint distribution, over signatures and exposures, is

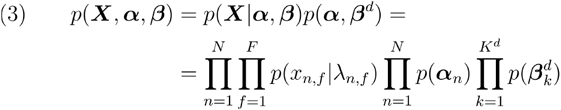

with *α*_*n*_ and 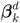 following a Dirichlet prior with a concentration that ensures sparsity among the de novo and catalogue signatures (Online Methods). Our approach precisely forces the model to sample from features with low density in the catalogue, ensuring that de novo signatures account for the signal unexplained by catalogue signatures. To optimise the number of de novo signatures, we minimise the Bayesian Information Criterion (BIC) computed using the maximum a posteriori estimates of the model parameters. Moreover, we implement a final heuristic to drop de novo signatures that are linear combinations of other de novo or catalogue signatures. This heuristic helps retain signals that differ from one another, also when considering complex combinations of signatures. Details on the overall formulation are in the Online Methods.

### Non-parametric clustering using multiple types of mutational signatures

Patterns of similar *α* across patients can unravel distinct molecular subtypes, suggesting that patients could be clustered by exposures of multiple types of signatures (i.e., SBS, DBS, etc.) as mutagenic processes might cause the emergence of different mutagenic patterns. Once signatures have been extracted with independent bNMF runs, BASCULE can use a Bayesian non-parametric model (Fig. 1D) to cluster *N* patients into *G* ≥ 1 groups with similar exposure patterns; to cluster *V* ≥ 1 type of signatures at once, our model uses a tensor-based approach that leverages a Dirichlet process [33], and automatically determines the number of clusters.

BASCULE represents the input as a tensor ***A*** with dimensions 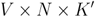, where 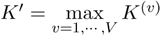 for *K*^*(v)*^ the number of columns (i.e., signatures) of the -th type of signature. A zero-padding strategy that flattens out the difference in the number of dimensions per signature type is required to assemble ***A***. Similarly to the bNMF step, the exposures are modelled with a Dirichlet, leading to a mixture likelihood

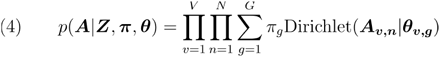

and the factored joint probability

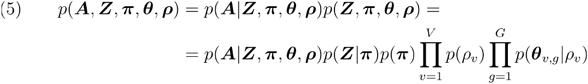

BASCULE defines the distribution over the mixture components and their weights as a Dirichlet Process *DP*(*η,H*) with concentration *η* and base distribution *H* such that is ***θ***_*g*_ sampled from a Dirichlet. In the implementation, we provide the model with an input value representing the maximum number of clusters, and use a stick-breaking construction to sample from the Dirichlet Process [34]. With this approach, the model automatically determines the optimal number of clusters based on the concentration *η*. After clustering, BASCULE merges clusters with similar centroids based on cosine similarity; this heuristic helps retain clusters with different exposures to one another. The derivations of the model are available as Online Methods.

### Synthetic simulations

We conducted realistic simulations by sampling from the generative model of BASCULE (Fig. 1C,D) and measured performance for various combinations of the number of patients *N*, the number of groups *G* (molecular subtypes), and mutational processes *K* (example dataset in Fig. 2A). Our datasets have 150, 500 or 1000 samples, 1, 3, or 6 groups, and incorporate up to 16 SBS COSMIC signatures. For each configuration, we generated 30 independent datasets. We further validated our model by generating more realistic datasets using the SigFitTest tool [35], a recent method generating mutation catalogues from real data. In these simulations, datasets were generated with 150, 500, or 1000 patients and 1, 3, or 6 tumor types, which we considered as groups. Furthermore, we inspected datasets with 100, 2000 and 50000 total mutations and simulated both whole-exome (WES) and whole-genome sequencing (WGS) data. The number of SBS COSMIC signatures per dataset was automatically selected by SigFitTest, with a maximum of 39.

**Fig. 2.**
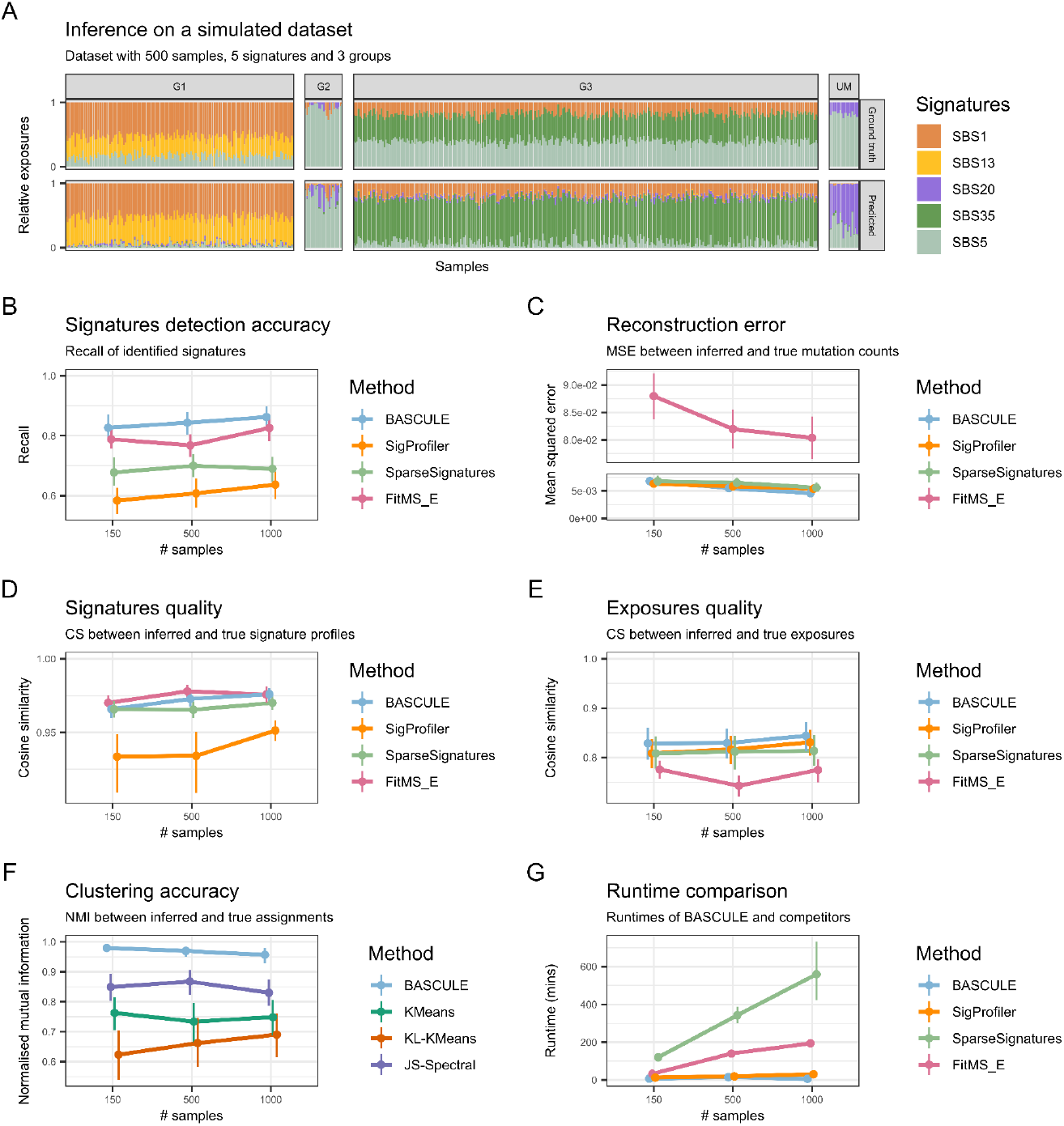
A. Example of inference on a simulated dataset. The generated dataset (top panel) consists of 500 samples, 3 groups, and 5 SBS signatures, including the profiles of COSMIC SBS13, SBS20, and SBS36 as de novo. The results (bottom panel) demonstrate that BASCULE successfully retrieves the signatures and learns similar exposures (cosine similarity=0.94). Moreover, BASCULE can group the samples in G1, G2 and G3 with high accuracy (NMI=0.93), with only a few unmatched samples (UM). B. Performance evaluation of BASCULE on synthetic datasets with an increasing number of input samples, compared to SigProfiler, SparseSignatures and FitMS extraction algorithm. Accuracy was assessed using recall, the number of true positives divided by the number of true positives plus the number of false negatives. C. Reconstruction error on the input mutation counts matrix. D. Quality of correctly retrieved signatures measured by cosine similarity between true and inferred parameters. E. Quality of exposures measured by cosine similarity between the full true and inferred parameters (Online Methods). F. Clustering accuracy was evaluated using the normalised mutual information, compared to K-means, K-means with Kullback-Leibler distance and Spectral clustering with Jensen-Shannon divergence of the exposure vectors. G. Runtime for SigProfiler, SparseSignatures and FitMS extraction algorithm compared to BASCULE during inference.

We compared the performance of the bNMF model to SigProfiler [36], the most widely used tool for de novo signature discovery, to SparseSignatures [24], a method that uses LASSO regularisation with a background signature, and to FitMS [23, 27], which performs signature extraction (FitMS_E) and fitting using constrained NMF with bootstrap-based robustness. Even if there are other tools for signature extraction, these tools represent a particular set of statistical approaches. Whereas SigProfiler is the historical method used to develop COSMIC, SparseSignatures and FitMS, like BASCULE, use prior biological information and distinct statistical approaches (lasso, bootstrap, and Bayesian inference) to achieve a parsimonious explanation of the data (Online Methods).

Results from model recall (Fig. 2B), i.e., the proportion of all actual positives classified correctly as positives, indicated that BASCULE can accurately identify the true number of signatures (Additional file 1: Fig. S1), in most settings better than competitors (Additional file 1: Fig. S2). Moreover, the quality of the deconvolution was high, as measured with the mean squared error (MSE) of the reconstructed mutation count matrix (Fig. 2C), and of the cosine similarity (CS) of the estimated signatures and exposures (Fig. 2D,E). In our tests, BASCULE, SigProfiler and SparseSignatures demonstrated the best counts reconstruction (BASCULE MSE 6.8e^-3^, SigProfiler MSE 6.3e^-3^, SparseSignatures MSE 6.9e^-3^, FitMS_E MSE 8.80e^-2^). All methods achieved high CS for signatures (BASCULE CS 0.97, SigProfiler CS 0.93, SparseSignatures CS 0.97, FitMS_E CS 0.97) and exposures (BASCULE CS 0.87, SigProfiler CS 0.79, SparseSignatures CS 0.81, FitMS_E CS 0.75). SigProfiler showed less precise signature estimation, and FitMS_E underperformed in exposure inference, whereas BASCULE and SparseSignatures showed consistent performance across metrics (Additional file 1: Fig. S2). This indicates that our model can reconstruct the signature profile and contribution to a sample. As expected, performance improved across all analyses and methods as the number of samples increased and the number of signatures decreased. Conversely, as the number of signatures increased, performance differences between methods decreased (Fig. S1 and S2; Online Methods).

Moreover, we evaluated the performance of BASCULE, the only method to cluster exposures across multiple signature types, against three alternative approaches that are generically applicable to the exposure vectors: a baseline k-means [37] algorithm using Euclidean distance (KMeans), a k-mean with Kullback-Leibler divergence (KL-KMeans) and Spectral Clustering [38] with Jensen-Shannon divergence (JS-Spectral) (Online Methods). Clustering performance was assessed using the normalised mutual information (NMI) between simulated groups and retrieved cluster labels (Fig. 2F). BASCULE Dirichlet-based clustering demonstrated superior performance than the other methods, achieving an NMI of 0.96 compared to 0.83 of JS-Spectral, 0.73 of KMeans and 0.62 of KL-KMeans and supporting the practical advantage of employing a signature-based model (Additional file 1: Fig. S3).

Finally, we measured the runtime of BASCULE on these synthetic tests and compared it to existing methods. We could perform this test at least for signature deconvolution (bNMF step of BASCULE), which is available in all competing tools (Fig. 2G and Additional file 1: Fig. S4). In this case, BASCULE proved to be faster than the other methods, which required 7 (BASCULE), 13 (SigProfiler), 32 (FitMS_E) and 120 (SparseSignatures) minutes in datasets with 150 samples, and 15 (BASCULE), 19 (SigProfiler), 140 (FitMS_E) and 343 (SparseSignatures) minutes for signature deconvolution of 500 samples. In particular, for large cohorts with 1000 samples, BASCULE was run on a GPU, resulting in a runtime of 6 minutes, compared to 30, 194 and 559 minutes for SigProfiler, FitMS_E and SparseSignatures. These results suggest that BASCULE can scale efficiently to analyse large cohorts.

Validation on SigFitTest datasets was more challenging than on those generated directly from the BASCULE generative model, resulting in a lower overall performance but similar qualitative behaviour across methods. All evaluated methods were particularly sensitive to datasets with a low number of mutations, consistent with previous observations [35]. Increasing the number of signatures, which increases the data complexity, also led to reduced performance. In contrast, sequencing type (WES and WGS) had minimal impact on the overall accuracy. Detailed results for this analysis are described in Online Methods and Fig. S5-S7.

### Molecular subgroups across three main human cancers

We used BASCULE to analyse WGS data recently released by [23], which carried out one of the most significant attempts to establish a catalogue of SBS and DBS signatures beyond COSMIC [29]. One of the critical advances of [23] *et al*. is to establish, using multi-step heuristics, which signatures are common across patients, as opposed to rare ones. Degasperi *et al*. used 12,222 WGS cancer samples collected through the Genomics England 100,000 Genomes Project [39] and available at Genomics England [40] to discover 40 SBS and 18 DBS mutational signatures in addition to COSMIC, for a total of 135 SBS and 180 DBS signatures in 19 tumour types. Additionally, Degasperi *et al*. compared their results with 18,640 samples from two external cohorts: the International Cancer Genome Consortium [41] and the Hartwig Medical Foundation [42]. In this paper, we used all data from Degasperi *et al*. in combination with the COSMIC and the Degasperi *et al*. catalogues, focusing on 6,923 samples from breast, lung, and colorectal cancers.

For each tumour type, we ran BASCULE with a minimal catalogue curated from the common signatures reported by Degasperi *et al*., using the COSMIC version if available (details in the Online Methods). In brief, we selected 10 SBS and 3 DBS signatures for breast, 11 SBS and 4 DBS for lung, and 13 SBS and 4 DBS for colorectal cancers. After bNMF for up to 25 de novo signatures, we used cosine similarity to map de novo signatures to the full COSMIC and Degaspari *et al*. catalogues (Additional file 1: Fig. S8); this ideally would allow us to detect less common signatures in our data. Ultimately, we performed clustering and used the centroid and patient exposures to derive meaningful insights from clustering assignments.

### Breast cancer patients (n = 2682)

From 2,682 breast cancer samples, we determined 23 signatures (13 de novo) and 5 clusters (Fig. 3A; Additional file 1: Fig. S9). Upon post-inference mapping (Fig. S8, S10), 10 out of 13 SBS and DBS de novo signatures were mapped to COSMIC, and two to the Degasperi *et al*. catalogue (Fig. 3B), with one flagged as rare in the original study.

**Fig. 3.**
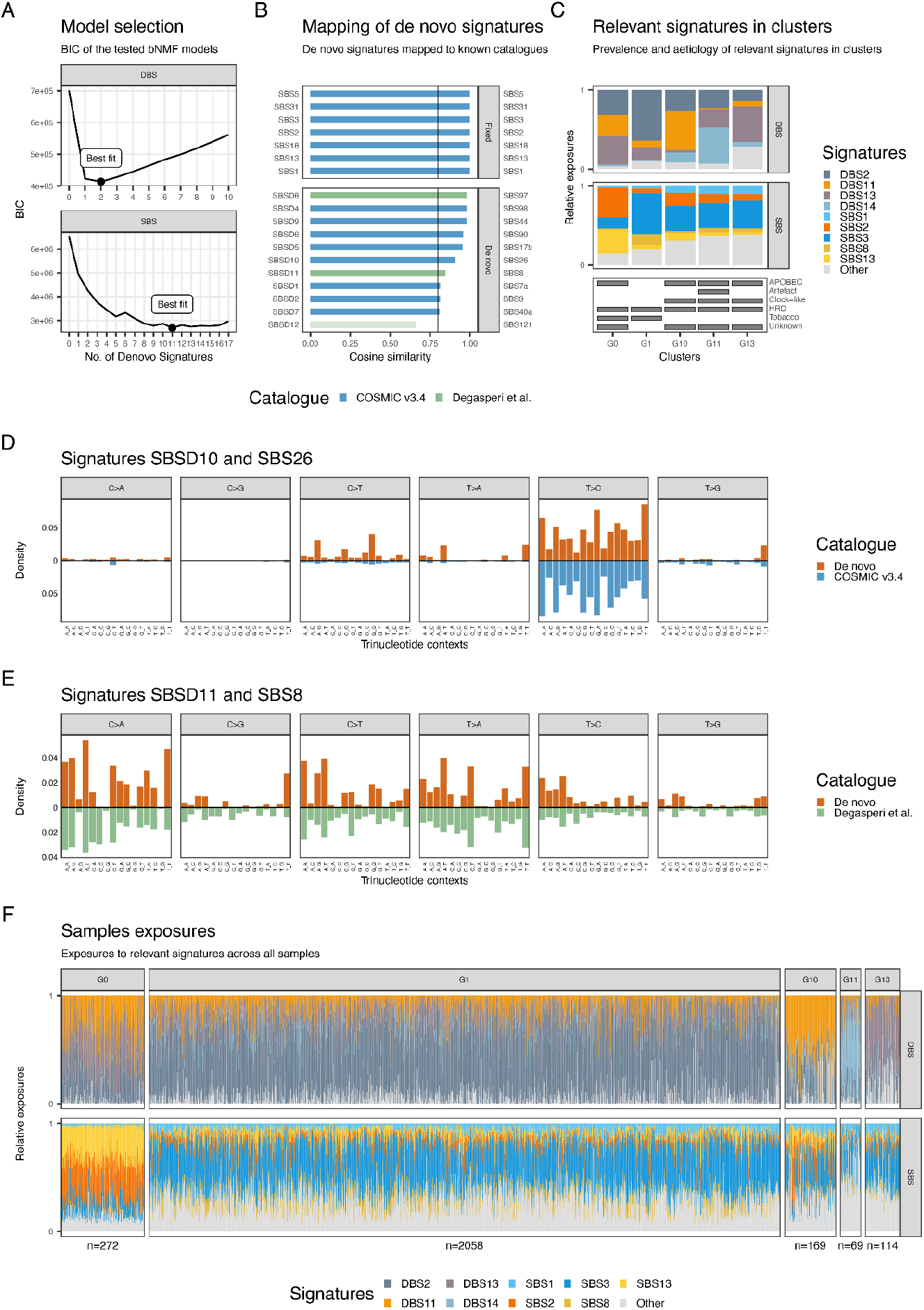
A. Optimal number of de novo SBS and DBS signatures based on BIC criteria, running BASCULE on 2,682 breast cancers with a starting catalogue of known signatures (SBS1, SBS2, SBS3, SBS5, SBS8, SBS13, SBS17, SBS18, SBS31, and SBS127 for SBS and DBS11, DBS13, and DBS20 for DBS) from COSMIC. B. Mapping for the inferred de novo SBS signatures to known catalogues. C. The exposure of the centroids for the clusters where only the most relevant signatures in each cluster are present. It includes clusters with more than 20 patients with their corresponding aetiology at the bottom. The full plot is accessible in Additional file 1: Fig. S9. D. De novo signature SBSD10 compared to the mapped signature SBS26 from COSMIC v3.4. E. De novo signature SBSD11 compared to the mapped signature SBS8 from Degasperi et al. F. Exposures of relevant signatures for clusters with more than 20 samples. We retained only those mutational signatures for which at least one patient cluster demonstrated a significant level of exposure and others labeled as “Other”.

We found two major biological interpretations for the 5 clusters, joining SBS and DBS signatures to explain all groups (Fig. 3C; Additional file 1: Fig. S11). These interpretations are facilitated by the potential alignment of some de novo signatures with already established known signatures (Fig. 3D,E). None of the clusters in breast cancer is featured with de novo signatures. Cluster G1 (n=2,058; Fig. 3F and Additional file 1: Fig. S11), the largest among all, showed a strong presence of SBS3, SBSD11 (mapped to SBS8 from Degasperi et al.), DBS2 and DBS13. SBS3 and DBS13 are caused by homologous recombination deficiency (HRD); DBS2, instead, is linked to smoking in some cancers but is often observed also in cancers not directly linked with tobacco exposure. Based on these prevalent signatures, cluster G1 can be associated with triple-negative breast cancers [43]. In cluster G0 (n=272), the dominant mutational signatures (SBS2 and SBS13, and evenly DBS11, DBS13 and DBS2) are associated with APOBEC activity, linking this group to HER2 breast cancers [44]. Clusters G10 (n=169), G11 (n=69), and G13 (n=114) share similar SBS patterns, including exposure to SBS1, SBS2, SBS3, SBSD11, and SBS13, pointing to a mix of APOBEC activity, HRD, and ageing-related mutational processes. BASCULE can differentiate these three clusters based on DBS exposure. In particular, G10 is strongly linked to DBS2 and DBS11, G11 primarily to DBS14 and G13 to DBS13 and DBS20. Therefore, G10 can be associated with APOBEC activity, G13 with HRD-related processes, and G11, the smallest cluster, with DBS14, a signature reported as artifactual in COSMIC. However, the presence of DBS13 in G11, suggests that this cluster is also linked to HRD. We remark that these clusters would not have been detectable by SBS patterns alone.

### Lung cancer patients (n = 1396)

From 1,396 lung cancer samples, we identified 7 distinct clusters characterised by 21 mutational signatures (9 de novo) (Fig. 4A; Additional file 1: Fig. S12). Following the mapping stage (Fig. 4B; Additional file 1: Fig. S8), 6 SBS and DBS de novo signatures were found in COSMIC and 3 in the Degasperi *et al*. study (one flagged as rare). As for breast cancer, we observed that by leveraging both SBS and DBS (Fig. 4C and Additional file 1: Fig. S12), as well as mapping some de novo signatures to known signatures (Fig. 4D,E), we uncover potential biological differences missing from SBS data alone. None of the clusters in lung cancer is featured with de novo signatures. The exposure patterns in lung cancer (Fig. 4F; Additional file 1: Fig. S13) seemed more complex than in breast cancer, with many clusters sharing a significant number of signatures and similar exposures.

**Fig. 4.**
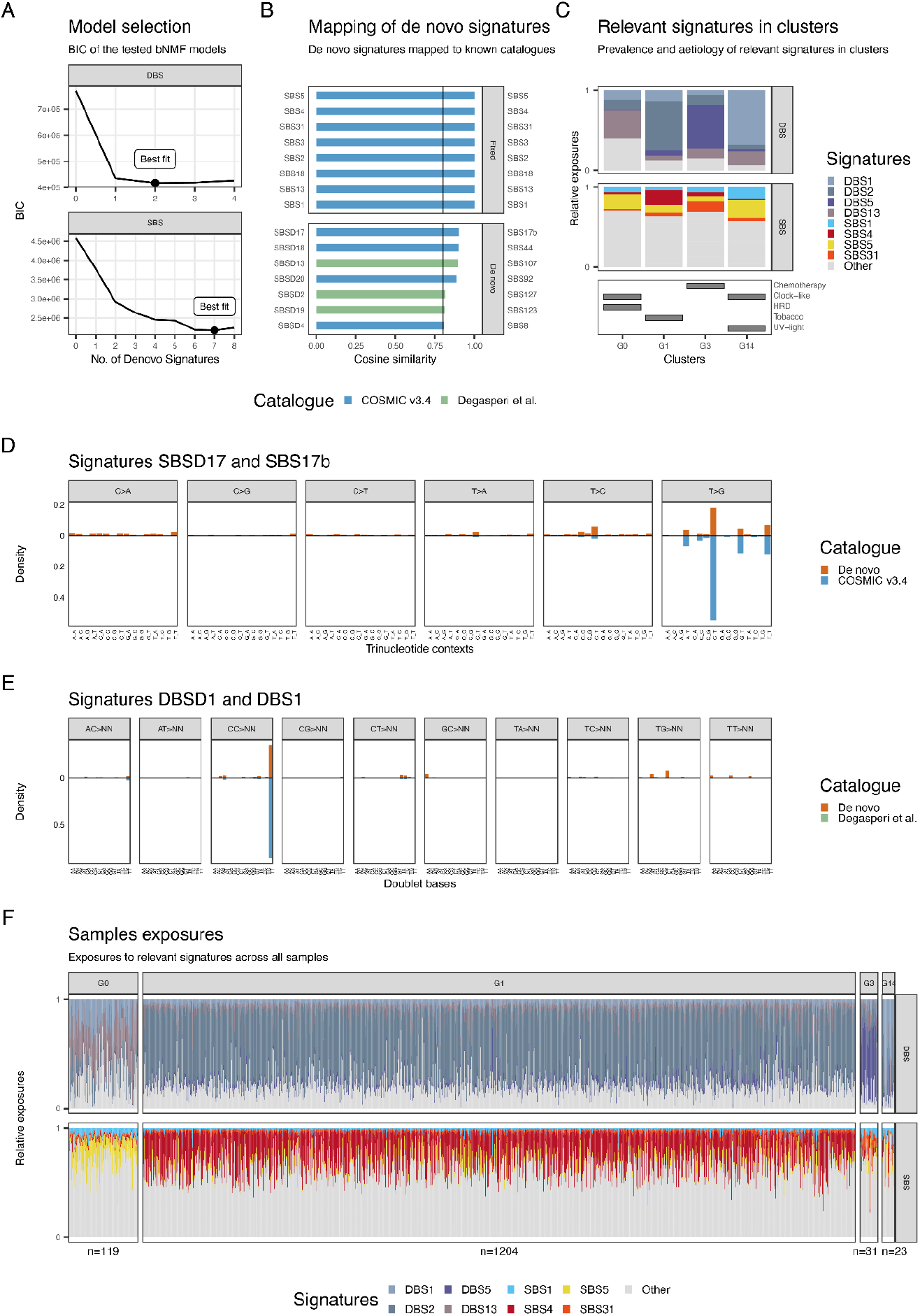
A. Optimal number of de novo SBS and DBS signatures based on BIC criteria, running BASCULE on 1396 lung cancers with a starting catalogue of known signatures (SBS1, SBS2, SBS3, SBS4, SBS5, SBS8, SBS13, SBS17, SBS18, SBS31, and SBS92 for SBS and DBS2, DBS5, DBS13, and DBS20 for DBS) from COSMIC. B. Mapping for the inferred de novo SBS signatures to known catalogues. C. The exposure of the centroids for the clusters where only the most relevant signatures in each cluster are present in the plot. It includes clusters with more than 20 patients with their corresponding aetiology at the bottom. The full plot is accessible at (Additional file 1: Fig. S12). D. De novo signature SBSD17 compared to the mapped signature SBS17b from COSMIC v3.4. E. De novo signature DBSD1 compared to the mapped signature DBS1 from COSMIC v3.4. F. Exposures of relevant signatures for clusters with more than 20 samples. We retained only those mutational signatures for which at least one patient cluster demonstrated a significant level of exposure and others labeled as “Other”.

Cluster G1 (n=1,204), the largest cluster, was associated with heavy smokers based on moderate exposure to SBS4, and pronounced exposure to DBS2 [45]. Another cluster, G3 (n=31), harboured SBS31 and DBS5, two signatures linked to chemotherapy exposure. Chemotherapy was also linked to cluster G13 (n=8), but this time because of SBS17b, presenting a unique pattern among the clusters. Both G3 and G13 contained, therefore, patients exposed to chemotherapy. Cluster G14 (n=23), compared to others, showed a slightly elevated presence of SBS5 and significant exposure to DBS1. As for breast cancers, the causative process for DBS1 remains unknown, whereas in other tumours, this signature can be linked with UV light exposure [46]. Finally, clusters G0 (n=119), G12 (n=4), and G8 (n=7) were similar in terms of SBS patterns but differed for DBS profiles; namely, G0 was characterised by DBS13, related to HRD, while G12 exhibited DBS6 and G8 DBS20, which have unknown aetiology (Online Methods).

### Colorectal cancer patients (n = 2845)

From n=2,845 colorectal cancer patients, BASCULE identified 32 signatures and 8 clusters of patients (Fig. 5A; Additional file 1: Fig. S14). Twenty signatures were SBS and DBS de novo, but 12 were mapped to COSMIC, and two to the Degasperi *et al*. study, both as rare signatures (Fig. 5B; Fig. S8, S15). Among our case studies, this dataset has the highest number of clusters and signatures (Fig. 5C) and mapping some de novo signatures to known signatures provides additional insights into the explanation of the clusters identified by our model (Fig. 5D,E). The remaining unmapped de novo signatures (Additional file 1: Fig. S15) did not feature in defining clusters.

**Fig. 5.**
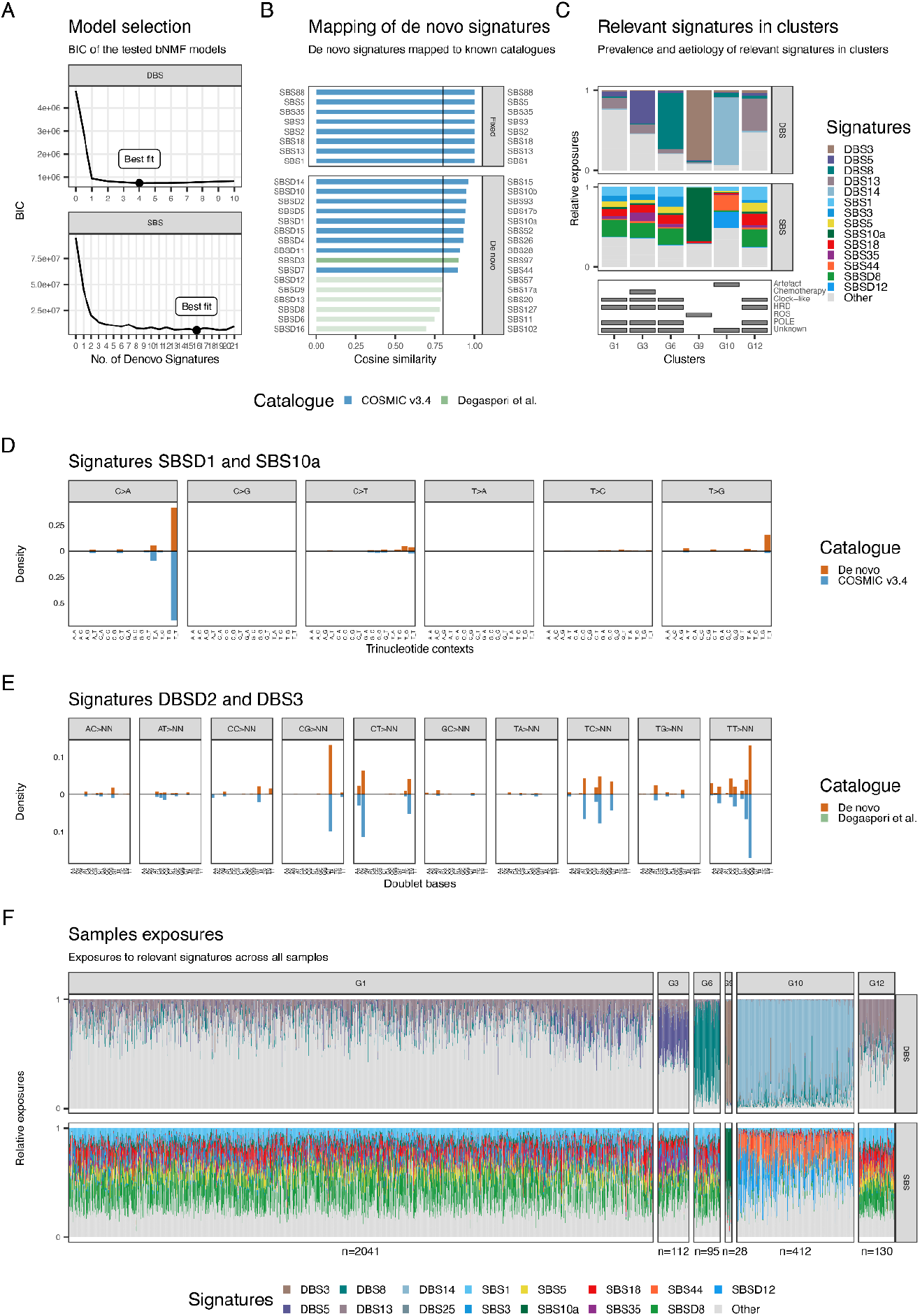
A. Optimal number of de novo SBS and DBS signatures based on BIC criteria, running BASCULE on 2845 colorectal cancers with a starting catalogue of known signatures (SBS1, SBS2, SBS3, SBS5, SBS8, SBS13, SBS17, SBS18, SBS35, SBS88, SBS93, SBS121, SBS157 for SBS and DBS2, DBS5, DBS13, DBS20 for DBS) from COSMIC. B. Mapping for the inferred de novo SBS signatures to known catalogues. C. The exposure of the centroids for the clusters where only the most relevant signatures in each cluster are present in the plot. It includes clusters with more than 20 patients with their corresponding aetiology at the bottom. The full plot is accessible at (Additional file 1: Fig. S14). D. De novo signature SBSD1 compared to the mapped signature SBS10a from COSMIC v3.4. E. De novo signature DBSD2 compared to the mapped signature DBS3 from COSMIC v3.4. F. Exposures of relevant signatures for clusters with more than 25 samples. We retained only those mutational signatures for which at least one patient cluster demonstrated a significant level of exposure and others labeled as “Other”.

At baseline (Additional file 1: Fig. S16), we could retrieve the canonical split among microsatellite stable (MSS) and unstable (MSI) colorectal cancers, including polymerase-eta (POLE) subtypes [47–49]. Clusters G1 (n=2,041), G3 (n=112), G6 (n=95), and G12 (n=130) displayed similar patterns of mutational signatures and exposures (Fig. 5F). These clusters prominently featured a BASCULE de novo SBSD8 signature. Quality control on all de novo signatures revealed that this signature is neither a linear combination of other signatures nor similar to any known signature (minimum tested cosine similarity 0.8). In addition, we found ageing-related signatures SBS1 and SBS5 alongside with, which are expected in most patients. Additional signatures were SBS44 (mismatch repair deficiency), SBS35 (chemotherapy), SBS3 (HRD), and SBS18 (reactive oxygen species). The tensor-based clustering approach distinguished these clusters by DBS signatures. Cluster G6 was characterised by hyper-exposure to the DBS8 signature, G3 by the unique DBS5 signature linked to chemotherapy, G12 by DBS13 associated with HRD, and G1 by the predominant DBS25 signature of unknown aetiology. Cluster G10 (n=412) reported exposure to the BASCULE de novo signatures SBSD12 and SBSD7. SBSD12 is mapped to SBS57 related to sequencing artefacts, and SBSD7 is mapped to signature SBS44 related to MSI status. In the DBS context, this cluster shows hyper-exposure to DBS14, which is considered artifactual in COSMIC. Cluster G9 (n=28) showed high exposure to SBS10a and DBS3, both indicative of the POLE status. Cluster G0 (n=23) featured the SBS18 signature and little of SBS3, two signatures linked with reactive oxygen species aetiology. In cluster G5 (n=4), the most relevant signature was SBS15, linked to mismatch repair deficiency, alongside notable exposure to the de novo SBSD7 signature identified by BASCULE which is mapped to SBS44 from COSMIC catalogue and related to MSI status (Online Method).

### The prognostic power of stratifications derived from signatures

BASCULE can be used to cluster patients based on mutational signatures, posing the question if this stratification generates molecular subgroups that can, at least in principle, become prognostic. Since a similar result was shown in [50], where they classified patients using their mutational signatures exposure patterns, we conducted survival analysis to assess the prognostic power of subgroups derived from samples with matched clinical information.

Precisely, we performed survival analysis using Kaplan-Meier estimation for overall survival trends, and multivariate Cox proportional hazards regression to evaluate if BASCULE cluster membership is an independent prognostic factor, after adjusting for clinical covariates such as age and gender. We limited this analysis to a subset of ICGC [51] samples – specifically, 259 skin, 343 pancreatic, and 315 esophageal tumor samples – with matched clinical data, and focused on SBS since DBS were very sparse (>98% zero entries; Online Methods).

In a study of n=259 skin cancer samples, we identified ten mutational signatures within two clusters, including the known established signatures like SBS1 (clock-like), SBS5 (clock-like) and SBS7 family (UV-light) (Fig. 6A and Additional file 1: Fig. S20). Kaplan-Meier survival analysis revealed distinct survival trends between the clusters (log-rank p=0.033; Fig. 6B). Cluster G11, which was enriched for SBS7 signatures, exhibited better survival compared to G1. In multivariate Cox regression (Fig. 6C), with G1 as reference due to its lowest survival outcomes, cluster G11 showed a trend toward a protective effect (n=190, hazard ratio=0.63, 95% CI: 0.39-1.02, p=0.059). This finding however was borderline significant, possibly because of the effect of age which significantly influenced the hazard: younger patients experienced a 44% reduction in risk relative to older individuals (n=114, hazard ratio=0.56, 95% CI: 0.40-0.80, p=0.001), whereas gender did not significantly affect the outcome.

**Fig. 6.**
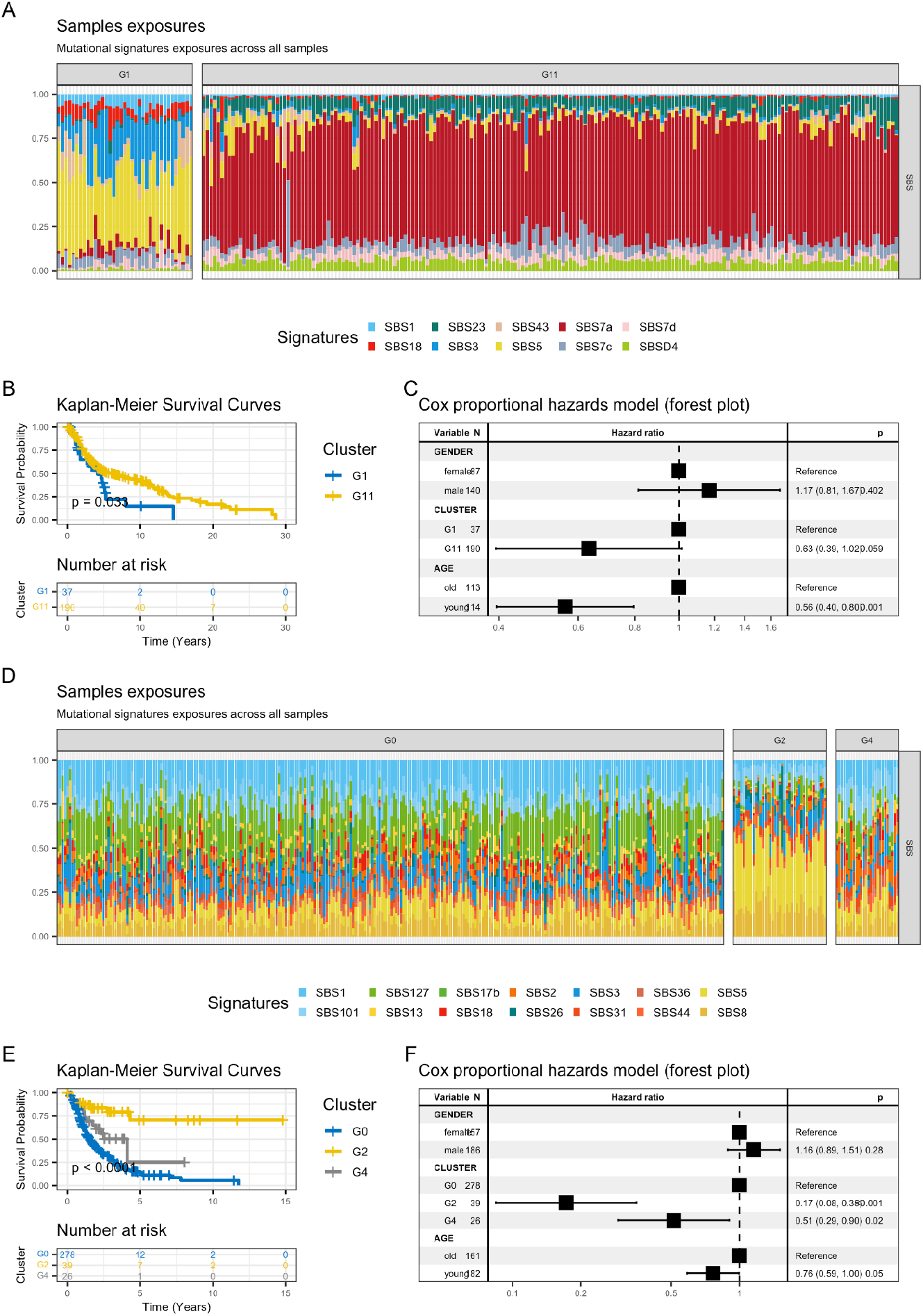
A. Exposure plot illustrating mutational signature profiles of skin cancer for clusters with more than 20 samples, highlighting the distribution of signatures across clusters G1 and G11. B. Kaplan-Meier survival plot depicting survival probabilities for clusters G1 and G11 over a 30-year period, with a risk table at the bottom. C. Forest plot, evaluating the significance of covariates (gender, cluster, and age) in the skin tumor dataset. D. Exposure plot illustrating mutational signature profiles of Pancreas cancer for clusters with more than 20 samples, highlighting the distribution of signatures across clusters G0, G2 and G4. E. Kaplan-Meier survival plot depicting survival probabilities for clusters G0, G2 and G4 over a 15-year period, with a risk table at the bottom. F. Forest plot, evaluating the significance of covariates (gender, cluster, and age) in the pancreas tumor dataset.

With a similar design we examined n=343 pancreatic tumor samples, identifying 14 mutational signatures in three distinct subgroups, all corresponding to predefined signatures in the COSMIC and catalogues from [23] (Fig. 6D and Additional file 1: Fig. S21). Among these, SBS1 and SBS5 (age related), SBS2 (APOBEC), SBS3 (homologous recombination deficiency) and SBS26 (mismatch repair deficiency) are well-known signatures. The analysis of survival curves (Fig. 6E) indicated a significant difference in survival outcome across clusters G0, G2 and G4 (log-rank p < 0.0001). Cluster G2, dominated by SBS5, had the highest survival, confirmed by Cox proportional hazard analysis (Fig. 6F) with cluster G0 as reference. G2 (n=39, hazard ratio=0.17, 95% CI: 0.08-0.35, p < 0.001) exhibited an 83% lower hazard than G0, and G4 (n=26, hazard ratio=0.51, 95% CI: 0.29-0.90, p=0.02) a 49% lower hazard than G0. In this tumour these results were both statistically significant, with age providing a protective effect – 24% lower hazard– in the younger (n=182, HR=0.76, 95% CI: 0.59-1.00, p=0.05) compared to the older group (reference). In this tumour, gender was associated with a hazard ratio of 1.16 (95% CI: 0.89-1.51, p=0.28), but did not affect outcome.

We also investigated esophagus cancer patients with a similar design, but our analysis did not reveal significant differences in survival trajectories. We report that analysis in the Online Methods.

## Discussion

Over the last few years, large-scale analyses revealed ubiquitous mutational signatures across distinct cancer types. For this reason, signatures have become a pillar concept of modern cancer genomics, thanks to the possibility of linking their activity with endogenous and exogenous processes with known aetiology [30, 52], and translation [17, 18, 53]. The field is quickly advancing toward developing signature catalogues that stratify human tumours into distinct molecular subtypes, with COSMIC standing as the leading example of this progress.

The feasibility of this grand challenge strongly depends on the strength of the statistical signals recorded in genomics data, and might lead to limitations ubiquitous to computational tools. Some assays, e.g., WES, might be underpowered unless in some specific scenarios of strong mutagenesis (e.g., smokers in lung cancers, defective DNA repair in breast cancer etc.). Some tumours might also be less prone to signature analysis, like for instance certain leukemia or childhood cancers (e.g., medulloblastoma or astrocytoma) that generally show very low mutation rate. Overall, the concept of signatures remains broad and flexible, and whereas a sample might not be rich of somatic point mutations allowing SBS analysis, it might be rich of aneuploidy (e.g., testicular germ cell tumors, liposarcoma, etc.) to allow the extraction of copy-number signatures.

BASCULE offers a generic approach, which works for any type of signature and that addresses two major limitations of existing frameworks [21]: i) the inability to incorporate established catalogues in the search for new signatures, and ii) the lack of clustering capabilities based on signature data. We desire to build on existing catalogues to create new biological knowledge incrementally, and we want to identify groups of patients with similar signatures to translate signatures into biomarkers.

In this paper we first benchmarked BASCULE against state-of-the-art methods [23, 24, 27, 36]. Our tests showed that BASCULE can accurately retrieve signatures and infer high-quality exposures and signatures across many cases, with the differences between tools that become less pronounced as the number of active signatures increases to saturate the statistical signals.

We then created a minimal version of two known catalogues to test these ideas on three widespread human cancers, leveraging a GPU-based implementation of the Bayesian models of BASCULE. From about 7000 WGS samples, our approach retrieved off catalogue signatures that were never shown to our tool, but that were present in the extended version of the input catalogues. Notably, some were flagged as rare using dedicated heuristics in [23], suggesting that BASCULE is sensitive enough to detect signatures even when involving few patients. On the same data, BASCULE clustering retrieved molecular subtypes that are common in breast, lung and colorectal cancers. Our model autonomously detected groups that may be associated with triple-negative and hormone receptor-positive breast cancers, microsatellite-stable and unstable colorectal cancers, and lung cancers associated with tobacco exposure. Notably, compared to alternative tools, BASCULE clustering integrated SBS and DBS signatures, proving that combined analysis enhances stratification power. Moreover, in three other tumor types with matched clinical data (skin, pancreatic and esophageal), SBS signatures analysis yielded stratifications that were often prognostic, as we assessed via Kaplan-Meier analysis and multivariate Cox regression.

## Conclusions

BASCULE is a Bayesian method to integrate multiple types of mutational signatures, and stratify patients based on signatures occurrence. In the future, this framework could be extended to include additional molecular signals, such as those deriving from methylation and gene expression, assays. At a more technical level, advanced matrix factorisation strategy could also be developed to correlate inferences across samples collected from the same patient (e.g., in multi-region or longitudinal studies).

Overall, a generalised framework for signature deconvolution might ultimately help linking the emergence of specific cancer phenotypes with the underlying mutagenic processes, facilitating experimental investigations while offering a principled method to expand established biological signature catalogues.

## Availability of data and materials

The open-source BASCULE R package is available under the GPL-3.0 license, and can be accessed at https://github.com/caravagnalab/bascule [54, 55].

The Nextflow [32] module of BASCULE will be available in the nf-core [56] official repository https://github.com/nf-core/modules [56].

The input datasets utilised in this manuscript, which were derived from the Genomics England (GEL), International Cancer Genome Consortium (ICGC), and Hartwig Medical Foundation (HMF) cohorts, are publicly accessible at [23]. The clinical data utilised for the survival analyses is publicly downloadable from the ARGO Data Platform (https://www.icgc-argo.org/).

Additionally, all source code used for the analysis and reproduction of all the results described in this manuscript is available at https://doi.org/10.5281/zenodo.17192805 [55].

These resources provide researchers with the tools and data to replicate this study.

## Acknowledgements

The authors acknowledge the AREA Science Park supercomputing platform ORFEO made available for conducting the research reported in this paper and the technical support of the Laboratory of Data Engineering staff. The authors acknowledge the ICSC for awarding this project access to the EuroHPC supercomputer LEONARDO, hosted by CINECA (Italy).

## Author information

### Contributions

EB, AS, RB, SM and GC conceptualised and developed the model. EB, AS, RB and SM implemented the tool. EB, AS and EV carried out simulations and analysed real data. EB, AS, DR, NC and GC interpreted results. EB, AS and GC drafted the manuscript, which all authors approved in final form. GC supervised and coordinated this project. All authors read and approved the final manuscript.

## Ethics declarations

### Ethics approval and consent to participate

Not applicable

### Consent for publication

Not applicable

### Peer review information

Yuanhua Huang and Veronique van den Berghe were the primary editors of this article and managed its editorial process and peer review in collaboration with the rest of the editorial team. The peer-review history is available in the online version of this article.

## Supplementary information

### Additional file 1: Supplementary figures

**Fig. S1**. BASCULE performance on synthetic datasets. **Fig. S2**. BASCULE performance comparison on synthetic datasets. **Fig. S3**. BASCULE clustering accuracy on synthetic datasets. **Fig. S4**. BASCULE runtime on synthetic datasets. **Fig. S5**. BASCULE performance comparison on realistic WGS datasets. **Fig. S6**. BASCULE performance comparison on realistic WES datasets. **Fig. S7**. BASCULE clustering accuracy on realistic datasets. **Fig. S8**. Unmapped de novo signatures mapping to Degasperi and COSMIC catalogues in breast and colorectal cancers analysis. **Fig. S9**. Clustering centroids in breast cancer analysis. **Fig. S10**. Unmapped de novo signature in breast cancer analysis. **Fig. S11**. Exposures in breast cancer analysis. **Fig. S12**. Clustering centroids in lung cancer analysis. **Fig. S13**. Exposures in lung cancer analysis. **Fig. S14**. Clustering centroids in colorectal cancer analysis. **Fig. S15**. Unmapped de novo signatures in colorectal cancer analysis. **Fig. S16**. Exposures in colorectal cancer analysis. **Fig. S17**. BASCULE and FitMS exposures in breast cancer analysis. **Fig. S18**. BASCULE and FitMS exposures in lung cancer analysis. **Fig. S19**. BASCULE and FitMS exposures in colorectal cancer analysis. **Fig. S20**. BASCULE analysis of skin cancer data. **Fig. S21**. BASCULE analysis of pancreas cancer data. **Fig. S22**. Differential exposure scores.

## Online Methods

### The BASCULE framework

We developed a method, BASCULE, that can extract a broad spectrum of mutational signatures in a dataset, while leveraging the existing knowledge established from previous studies, expressed as a reference catalogue. The model will use Bayesian inference to search for de novo signatures that are statistically distinct from the reference ones, allowing to augment a catalogue with minimum overlap among signatures, the number of which is optimised by a likelihood criterion.

### Bayesian Non-negative Matrix Factorization

Signature deconvolution is carried out via Bayesian non-negative matrix factorization (bNMF), starting from a data matrix X with *N* rows (patients) and *F* columns (features). The entries of are non-negative values for the observed counts, and *F* depends on the type of signatures we wish to extract. BASCULE supports signatures from the simplest SBS, to DBS, ID and CN. For instance, when dealing with SBS and DBS, the features will be 96 and 78 strand-agnostic substitution contexts, respectively. In the general NMF step,*X* is factorised as

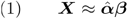

Here, *β* is the *K* matrix of signature profiles, while the matrix of signature activities 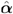 indicates, for each sample *n*, the number of mutations attributed to signature *k*, such that 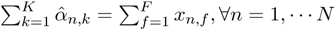. The bNMF model introduced here estimates the normalised exposure matrix, denoted as *α*, computed as the ratio of the signature activity matrix 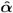 by the total number of mutations observed in each patient.

One key feature of BASCULE is its ability to embed prior knowledge of known signatures into the model. To this extent, the matrices *α* and *β* can be defined as follows

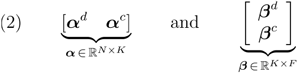

where the matrix *β* is defined as the concatenation of de novo signatures (*β* ^*d*^) to the reference catalogue (*β* ^*c*^), and the matrix *α* as the concatenation of exposures to de novo (*α* ^*c*^) and reference (*β* ^*c*^) signatures.

Assuming *K* signatures, we define *K*^*d*^ and *K*^*c*^ as the number of de novo and reference signatures, respectively. Hence, the total number of signatures is given by 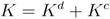. is decomposed in a *K* ×*F* matrix of signature profiles *β*, and a matrix *N* ×*K* of exposures α, such that

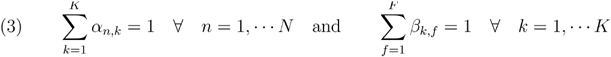

The non-negativity constraint is such that all entries of *α* and *β* are positive. To extend to model to be Bayesian, we use a Poisson likelihood

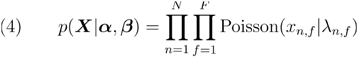

assuming the mutation classes to be independent, with the Poisson rate defined as the reconstructed mutation counts matrix 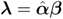.

The matrix *β* entries learnt during the inference are those regarding *β*^*d*^, while *β*^*d*^ is held fixed to reflect the reference. If *K*^*d*^ is set to 0, the model solely infers the latent exposures for the input catalogue of reference signatures. Otherwise, the model additionally learns de novo signature profiles *β*^*d*^ and their respective exposures.

The bNMF joint distribution can be factored as follows:

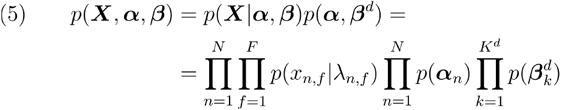

with priors

- 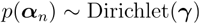
- 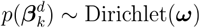

Since *α* and *β* are probability vectors, their priors are Dirichlet distributions with concentration γ (*K*-dimensional) and ω (*F*-dimensional). The value of ω can be specified as input to assign greater weight to particular signatures, such as tumour-specific ones. If no prior information is available, ω is set to a vector of ones. The value of ω is computed to maximise the distance between each de novo and reference signature. We construct a vector with low concentration in features that are highly represented in the reference catalogue by summing feature-specific contributions across reference signatures and then inverting and scaling these cumulative values. This approach forces the model to sample from contexts with low density in the reference, ensuring that de novo signatures will be used to account for the signal that remains unexplained by reference signatures.

The Maximum A Posteriori (MAP) for each latent parameter is estimated via Stochastic Variational Inference (SVI) in Pyro [31]. The model is parametric, and the number of signatures is required as input. The method selects the optimal number of de novo signatures as the one minimising the Bayesian Information Criterion (BIC), defined as 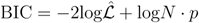. Here, *N* denotes the number of samples in the input data,*p* represents the model complexity, i.e., the number of inferred parameters, and 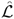 is the likelihood function (3) calculated using the values of the parameters.

In the current formulation of the method, we use a two-steps inference process. First, we run the model to infer the exposures of reference signatures, without introducing any de novo signatures, i.e., with 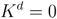, and filter out signatures from the catalogue with low exposures, resulting in a reduced catalogue 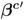. Specifically, the retained reference signatures, denoted as 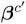, are those with normalised exposure above a predefined threshold (0.2 by default) in at least one sample. In the second step, we include the significant reference signatures 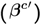 and infer any new de novo one (*β*^*d*^). In both steps of inference, the signature profiles in the reference catalogue remain fixed, the only change is in the number of considered signatures. This approach minimises noise from the reference catalogue and maximises the signal attributed to already validated signatures.

### Post-fit heuristics

After inferring de novo signatures, the method can further evaluate their similarity relative to linear combinations of other signatures. This helps determine if one signature is actually a combination of other known signatures (or other de novo signatures). Specifically, we decompose each de novo signature profile into a linear combination of the remaining ones, as

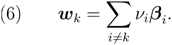

If the vector *w*_*k*_ closely matches – based on a cutoff on the cosine similarity – the original signature *β*_*k*_, the method will remove the de novo signature and redistribute its exposures according to the weights ν of this linear combination. This process reduces the number of signatures, enhancing the sparsity of the final results. We discuss this heuristic in the application to real data, together with specific parameters we used to refine our fits.

### Non-parametric Dirichlet mixture model

BASCULE can perform deconvolution of multiple signature types and cluster a cohort of *N* samples based on these multiple signals. Here, we are clustering a tensor *V* of exposure matrices into latent groups through a Dirichlet Process Dirichlet Mixture Model.

Assume to have *V* exposure matrices 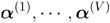 of *V* distinct types of signatures (i.e., SBS, DBS, IDS, etc.). Each matrix *α* has *N* rows (i.e., samples) and 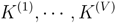 columns (i.e., signatures). To cluster the samples based on multiple signature types, we embed the *α* matrices in a tensor *A* of dimension 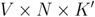. Since the deconvolution of distinct signature types might result in different optimal numbers of signatures, we need to standardise the dimensions to embed the exposure matrices into a tensor. Here, we define 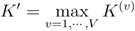 and perform zero-padding in the columns of *α*^(*v*)^ if 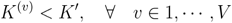.

Similarly to the bNMF step, since the exposures can be modelled with a Dirichlet, the overall mixture likelihood can be written as

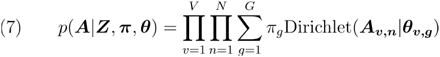

and the joint probability can be factored as

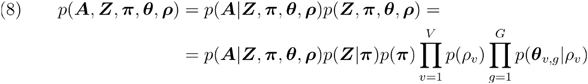

with

- *P(θ*_*v,g*_) ∽Dirichlet(ϕ_*v,g*_ · ρ_*v*_)
- *P (*ρ_*v*_) ∽ Gamma (*c,d*).

The model defines a distribution over mixture components and weights as a Dirichlet Process *DP*(η,*H*)with concentration η and base distribution *H*. We adopt the stick-breaking formulation of the Dirichlet Process to automatically determine the optimal number of clusters, up to a fixed maximum value *G*. More specifically, the model takes as input a value for the maximum number of clusters, and, after the inference has completed, clusters with no data points assigned are discarded. ***θ***_*v,g*_ is a *K* -dimensional vector reporting the component-specific exposure centroids for each signature. This is distributed as a Dirichlet whose concentration depends on ***ϕ***_*v,g*_, a *K*-dimensional concentration vector scaled by ***ρ***_*v*_. In the model, ***ϕ***_*v,g*_ might account for prior information on each retrieved signature and is set to centroids estimated with K-means. The variable ***ρ***_*v*_ is a signature type-specific scaling factor used to lower the variance of the Dirichlet distribution and to allow sparsity in the centroids. The value ***ρ*** of is computed as described in the next paragraph.

The inference has been carried out via Stochastic Variational Inference in Pyro [31]. The parameters were initialised with K-means to determine the component centroids (*θ*). In the model, the centroids *θ* are distributed as a Dirichlet. One important property of the Dirichlet distribution is that the concentration parameter influences the variance of the generated samples. More specifically, low concentration values lead to high variance, and high concentration values result in low variance. To address this, we introduced a scaling factor (*ρ*) to reduce the variance of the *θ* samples. The goal of *ρ* is to adjust the distribution of generated samples so that it better aligns with the observed data. An appropriate value for *ρ* is determined as follows. After running K-means to initialise the parameters, we calculated an upper bound for the empirical variance of exposures for each cluster and signature type *v*, to grasp the natural variation in the exposure values. We then chose the value of *ρ* to ensure that samples drawn from the Dirichlet prior centred around *θ* prior would approximately match the observed variance. This approach ensures that the generated exposure distributions closely match the observed ones, leading to improved convergence of the model fitting to reliable estimates of the centroids.

The model infers the maximum a posteriori (MAP) set of parameters and assigns each data point to the cluster maximising the posterior distribution.

#### Post-fit heuristics

After inferring the latent clustering assignments, the method can evaluate whether some clusters should be merged based on the inferred centroids (). Specifically, the method iteratively calculates the cosine similarity between the centroids. If this similarity exceeds a custom threshold, the clusters are merged, and the centroids are updated. This process continues until all clusters are distinct according to the defined similarity measure. We discuss this heuristic in the application to real data, together with specific parameters we used to refine our fits.

### Simulated data

We evaluated BASCULE’s performance by examining the error in reconstructing mutation counts, the accuracy of signature retrieval, the quality of signature exposures and the accuracy of clustering. To do this, we generated two batches of simulated datasets.

The first one is composed of synthetic datasets of increasing complexity sampled from the BASCULE generative bNMF model. Specifically, we considered cohorts of 150, 500 and 1000 samples, composed of 1, 3 or 6 groups. As the number of groups increases, the number of signatures (SBS and DBS) also increases, ranging from 1 to 13. In all fits, certain signatures are shared among all clusters. In contrast, others are either private to a single cluster or shared by a specific combination of clusters, enabling the unique identification of each group of patients. For each configuration, we generated 30 datasets. Each fit has been run with SBS1, SBS5, DBS3 and DBS5 as reference catalogues, testing five different de novo signature values around the true one.

We then tested the performance of BASCULE and competing NMF methods on a batch of simulated datasets, generated using the “generate_synthetic_catalogs” function from the “SigFitTest” Python package [35]. This tool creates realistic synthetic mutation catalogues with known signatures and exposures collected from COSMIC. We generated datasets of 150, 500 and 1000 samples, each composed of 1, 3 and 6 tumour types. For each combination of number of samples and groups, we further simulated datasets with 100, 2000 and 50000 mutations per sample, using both Whole Exome and Whole Genome Sequencing. The number of signatures has been automatically selected by the method and spans a wider range with respect to the batch of synthetic datasets, up to 39 active signatures. For each configuration, we generated 30 datasets.

#### Performance metrics

We evaluated BASCULE and competitors NMF performance based on four criteria: i) the ability to identify the correct set of signatures, ii) the ability to correctly reconstruct the input mutation catalogues iii) the quality of the reconstructed signatures and iv) the quality of the inferred exposures. We further compared BASCULE’s clustering model with competitors based on v) the quality of clustering assignments.

After de novo signatures deconvolution on the input datasets, each tool produced a set of estimated parameters: a combination of de novo and known signatures, along with the corresponding exposures. We matched the identified de novo signatures to the true ones using cosine similarity, applying a threshold of 0.8. That is, a signature was classified as a true positive (TP) if its cosine similarity with a true signature exceeded 0.8, as a false negative (FN) if it was not retrieved by the method and as a false positive (FP) if it was identified by the method without being present in the ground truth.

We assessed the similarity between the retrieved de novo signatures and the simulated ones using recall and precision metrics. Recall was computed as the ratio of TP to the sum of TP and FN, representing the proportion of correctly identified signatures out of all true ones. Precision was calculated as the ratio of TP to the sum of TP and FP, indicating the proportion of correct de novo signatures among those retrieved. To evaluate the quality of mutation catalogue reconstruction, and signature and exposure quality, we used Mean Squared Error (MSE) and Cosine Similarity (CS). These are standard metrics in the field [21].

To evaluate the quality of mutation catalogue reconstruction, we computed the MSE between the true input mutation count matrix ***X*** and the reconstructed catalogue, expressed as 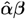, with 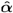 and β derived from the estimated exposures and signatures. We note that in FitMS_E, a subset of the mutation counts not attributable to mutational signatures is labeled as “Unassigned”. These “Unassigned” exposures were excluded when computing the MSE.

We assessed signature quality as the CS between true and inferred signatures classified as TPs. Exposure quality was evaluated through the CS between true and estimated exposure matrices. We first computed the CS for exposures corresponding to matched TP signatures. Then, we calculated the CS between the full true and inferred exposures matrices. To enable this comparison, we performed zero padding on the columns of the exposure matrices to include unmatched signatures. This allowed us to assess whether the FN and FP signatures had high activity in the samples. In the case of FitMS_E, we treated the “Unassigned” field as a FP signature.

We evaluated clustering performance by measuring the precision of reconstructing the correct grouping of patients, using the Adjusted Rand Index (ARI) and Normalised Mutual Information (NMI) between the true and inferred assignment vectors. The ARI compares how well the clusters predicted by a model match with the true clusters (ground truth), adjusting for the chance of random clustering, with a score ranging from -1 (completely incorrect clustering) to 1 (perfect clustering). NMI measures how much information is shared between the predicted clusters and the true clusters. The values range from 0 to 1, where 0 means the clustering assignments are independent, and 1 means identical (it is normalised to ensure comparability across different datasets, regardless of the size or number of clusters).

#### BASCULE performance on synthetic datasets

We evaluated BASCULE performance for the bNMF and clustering models. We first assessed, in terms of recall and precision, the similarity among the number of retrieved de novo signatures and the simulated one (Additional file 1: Fig. S1A,B). The recall results demonstrate that BASCULE effectively retrieves a precise number of de novo signatures, with only a few signatures missed in the analysis. The precision results indicate that only a few signatures were incorrectly identified. As expected, in both analyses, increasing the number of signatures results in a lower accuracy, whereas increasing the number of samples, thus providing a stronger signal for the signatures, improves the accuracy. Additionally, the performance is higher when working with SBS, which could be attributed to the greater sparsity of DBS signatures than SBS ones. Precision and recall showed that some groups have more inferred signatures than present in the ground truth, though these additional signatures typically have minimal exposure. This occurs because NMF was applied jointly across all groups to capture shared and group-specific patterns. Consequently, signatures with low or near-zero contributions may appear where they are not biologically meaningful. Such negligible exposures should not be interpreted as active signature presence.

We then assessed the quality of the mutation counts reconstruction, as well as the signatures and exposure learning. The input mutation counts matrix was reconstructed with high accuracy (Additional file 1: Fig. S1C), resulting in low values of MSE. The CS computed between true and estimated signatures (Additional file 1: Fig. S1D) and between true and estimated exposures (Additional file 1: Fig. S1E) shows great concordance between our results and the simulated datasets. Similar to recall and precision analysis, increasing the number of signatures adds complexity to the input data, resulting in reduced quality. However, increasing the number of samples leads to an improvement in the quality of the fit results.

Finally, we assessed the performance of the clustering model (Additional file 1: Fig. S1F). In our tests both metrics indicate that BASCULE performs well in accurately retrieving the correct sample assignments, with accuracy decreasing as the number of signatures increases and improving as the number of samples increases.

#### Comparison to other methods on synthetic datasets

Simulated counts were also used to compare BASCULE to three other methods for de novo mutational signature extraction: SigProfiler, SparseSignatures and FitMS_E. SigProfiler and SparseSignatures were run on 10 CPU cores of an AMD EPYC 7H12 computer, performing NMF with 100 repetitions for SigProfiler and 10 repetitions for SparseSignatures. FitMS was run to perform signatures extraction (FitMS_E) on 4 CPU cores of an Intel Xeon Gold 6140 computer, using the bootstrap signature fit approach with 10 replicates.

We compared the performance of the four methods to perform de novo signature extraction on SBS through the same metrics described in the previous paragraphs. BASCULE outperformed the other methods in identifying the correct number of signatures (Additional file 1: Fig. S2A,B), when the number of signatures was low, whereas all methods show lower values of recall and precision when increasing the number of signatures, and the difference in the methods performance decreases. Notably, BASCULE, SigProfiler and SparseSignatures showed low reconstruction error for the input datasets (Additional file 1: Fig. S2C), while FitMS_E showed a significantly higher MSE, probably due to the mutation counts labeled as “Unassigned” which correspond to a large proportion of the data.

For the correctly retrieved signatures, we assessed their quality based on cosine similarity (CS) (Additional file 1: Fig. S2D). BASCULE achieved higher CS scores than SigProfiler and SparseSignatures when the number of signatures was low, but all methods performed similarly as the number of signatures increased.

We also calculated the CS for the exposures of matched signatures (Additional file 1: Fig. S2E), where FitMS_E performance was lower than competitors with low number of signatures, and BASCULE’s performance decreased relative to other methods as dataset complexity increased. Finally, we computed the CS of exposures for matched and unmatched signatures, considering zero exposures for false negatives (FN) and false positives (FP) in the inferred and true exposures. Even in this case, FitMS_E showed a low similarity compared to competitors with low number of signatures. However, as dataset complexity increased, the CS of all methods became comparable, indicating that FN and FP are not necessarily signatures with low exposure levels.

We compared the clustering accuracy of BASCULE with k-means, KL-KMeans and JS-Spectral clustering methods performed on the exposures estimated by BASCULE (Additional file 1: Fig. S3). BASCULE outperformed all methods in all settings, especially when the number of signatures increased.

We finally compared the time required to fit each dataset for the three methods (Additional file 1: Fig. S4). BASCULE, implemented in Pyro, uses Stochastic Variational Inference (SVI), which enables faster inference compared to standard NMF methods. Additionally, BASCULE supports GPU-based inference, further accelerating computationally intensive calculations.

#### Comparison to other methods on realistic datasets

We compared BASCULE to the other methods using the batch of realistic datasets generated with SigFitTest [35] as explained in the above paragraphs. BASCULE and the competing methods were executed with the same arguments and HPC settings, and evaluated using the same metrics (Additional file 1: Fig. S5-S7). As expected, these datasets proved to be more challenging for all methods, which showed an overall lower performance. All methods were affected by the number of mutations, with performance particularly low for datasets with 100 mutations, as also noted in [35]. Similarly, an increase in the number of signatures, which increases dataset complexity, led to a decrease in overall performance across all methods. In contrast, the type of sequencing, i.e., whole exome (WES) or genome (WGS) sequencing, did not significantly impact the performance of the inference in any method or evaluated metric.

FitMS_E outperformed competing methods in terms of recall, demonstrating its ability to identify most of the correct signatures. However, its lower precision, particularly in datasets with few mutations, indicates a higher rate of FPs compared to other methods. BASCULE and SigProfiler showed similar performance in retrieving the correct set of signatures, with BASCULE better at avoiding FNs and SigProfiler more precise in signatures identification. SparseSignatures showed lower recall values but precision values comparable to other methods.

Overall, the reconstruction error is comparable across all methods, although BASCULE and SigProfiler demonstrated better performance in reconstructing the mutation catalogue when the number of mutations increases.

The correctly retrieved signatures (TPs) showed high cosine similarity with the true ones, with SparseSignatures displaying lower values compared to other methods.

The quality of estimated exposures for TP signatures improved with the number of mutations and remained consistent across methods. However, including FP and FN signatures in the evaluation led to reduced performance across all methods.

Finally, we compared the clustering accuracy of BASCULE compared to KMeans, KL-KMeans and JS-Spectral clustering methods performed on the exposures estimated by BASCULE (Additional file 1: Fig. S7). We used the information of the known tumour types provided by SigFitTest [35] as ground truth. However, this might not be entirely indicative of the actual number of groups in the datasets, as multiple tumour subtypes might be present and characterised by distinct signatures. The performance of all methods is quite variable, but BASCULE showed values of NMI with no significant differences.

#### SigProfiler

Version 1.1.21 of SigProfiler was run with the default values for the stability thresholds, using a minimum of 1000 and a maximum of 10000 NMF iterations, with a value of 1000 iterations for the convergence test. All other parameters were set to the default values.

#### SparseSignatures

Version 2.10.0 of SparseSignatures was run with the “normalize_counts” parameter set to TRUE, a value of 20 iterations and 10000 maximum lasso iterations. We used the “background2” signature, which is derived from COSMIC SBS5, as the background. For the cross-validation parameters, we used 5 entries, 10 repetitions, and a percentage of 0.01 entries to be masked by zeros. For sparsification, we tested values of 0 and 0.05 for α and values of 0, 0.01, 0.05 and 0.1 for β.

#### FitMS

Version 2.4.4 of the signature.tools.lib R package was used. In particular, we used the “SignatureExtraction” function to perform signatures extraction, setting the arguments “nboots” and “nrepeats” to 10 and 20, respectively, and providing a matrix of fixed signatures SBS1 and SBS5.

#### Clustering methods

We compared the BASCULE clustering Dirichlet Process model with three clustering methods: standard k-means with Euclidean distance, k-means with Kullback-Leibler distance (KL-KMeans), and spectral clustering with Jensen-Shannon divergence (JS-Spectral). For BASCULE, Model selection is performed automatically, whereas for the other methods, we use the Gap Statistic [57] to determine the optimal number of clusters.

For the standard k-means algorithm, we used the “kmeans” function from the R package “stats” [58], and performed model selection to determine the optimal number of clusters using the “clusGap” function from the R package “cluster” [59], executed with 10 bootstrap samples.

For the KL-KMeans method, we performed clustering using the “kcca” function from the R package “flexclust” [60], with a custom implementation of the KL distance employing the “kl.dist” function from the “seewave” R package [61]. Model selection was carried out using a custom gap statistic implementation adapted to the KL distance, with 10 bootstrap samples.

For the JS-Spectral method, we performed clustering using the “specc” function from the “kernlab” R package [62], with a custom implementation of the JS distance using the “dist” function from the “proxy” R package [63]. Model selection was similarly performed using a custom gap statistic implementation adapted to the JS distance, also ran with 10 bootstrap samples.

### Patient data analysis

#### Construction of the starting catalogue

We curated a simple version of a catalogue by combining our current understanding of mutational processes for SBS and DBS – the two types of mutational signatures for which we had data from [23]. The aim was to use this catalogue as input for our tool. To assemble it, we selected the common signatures from each tumour type as documented by Degasperi et. al. Subsequently if these particular signatures were also found in COSMIC, the corresponding COSMIC version of the signature was adopted for our input reference catalogue. In the final step, we enhance the signatures so that each one accurately reflects a distinct pattern of mutational signatures. This is achieved by eliminating signals from 96 contexts whose values fall below a specified threshold.

#### Input data

To perform the large-scale analysis discussed in the Main Text, we collected data from supplementary materials by Degasperi et al. Initially, we identified the organ-specific common signatures from Tables 9 and 10. Subsequently, we located their corresponding reference signatures in Tables 21 and 22, utilising the conversion matrix provided in Tables 25 and 26.

#### Fitting off-catalogue signatures

In the initial stage, we ran our tool, searching for a maximum of 25 de novo signatures (beyond the input catalogue). The model underwent the learning phase, consisting of 3000 iterations and a learning rate set at 0.005, and the most suitable number of de novo signatures was determined by evaluating the BIC score.

After inferring the latent variables, including the mutational signatures and exposure matrices, we applied a refinement step to the de novo signatures. This process aimed to identify and remove any de novo signatures that could be adequately explained by a linear combination of the other de novo and input catalogue signatures. The refinement method attempted to reconstruct each de novo signature using the coefficients of the corresponding linear combination. If the cosine similarity between the reconstructed signature and the original de novo signature exceeded 0.9 - an arbitrary threshold denoting very high similarity - the de novo signature was removed, and its exposure was redistributed among the signatures in the linear combination, weighted by their respective coefficients. By applying this refinement step, we ensured that the final set of de novo signatures represented mutational processes that could not be sufficiently captured by combining the existing signatures, either from the input catalogue or the newly identified de novo signatures. This approach helped to minimise redundancy and improve the interpretability of the mutational signature analysis.

#### Clustering patients by signature exposures

After this step, we ran the Dirichlet Process clustering. To optimise those results, we examined the clusters of signatures and merged those that exhibited some similarity. This process was conducted iteratively, merging clusters based on their similarity with their cluster centroids. The merging process continued until the cosine similarity between all pairs of clusters fell below 0.8, a predetermined cutoff value that denoted low similarity.

To interpret our clusters, we additionally assessed whether our de novo signatures could be accounted for by any signatures from the COSMIC database and the catalogue of Degaspari et al. To achieve this, we calculated the cosine similarity between all possible pairs of de novo signatures versus signatures from both extended catalogues. If the similarity exceeded 0.9, we replaced the de novo signature with the corresponding known signature from these catalogues, as discussed in the Main Text.

In this analysis phase, we examined the clusters of signatures and merged those that exhibited similarity. This process was conducted iteratively, merging clusters based on the similarity of their centroids. The merging process continued until the cosine similarity between all pairs of clusters fell below a predetermined cutoff value. Additionally, we assessed whether our de novo signatures could be accounted for by any signature known in COSMIC or the catalogue by Degaspari et al. To achieve this, we calculated the cosine similarity between all possible pairs of de novo signatures and the target catalogues, and, if the similarity exceeded 0.9, we replaced our de novo signature with the corresponding known one.

#### Breast cancer analysis

We analysed 2,682 breast tumour samples from the GEL, ICGC, and HMF cohorts. For the SBS context, we selected signatures SBS1, SBS2, SBS3, SBS5, SBS8, SBS13, SBS17, SBS18, SBS31, and SBS127, and for the DBS context, we selected signatures DBS11, DBS13, and DBS20, collectively forming the reference catalogue for breast tumour data.

The bNMF analysis identified 19 SBS (12 de novo) and 5 DBS (2 de novo) signatures. After a refinement step, we identified 18 SBS (11 de novo) and 5 DBS (2 de novo) signatures. The refinement process revealed that the de novo signature SBSD3 could be explained by the input catalogue signature SBS1, with a cosine similarity of 0.99 between the reconstructed and original signatures. As a result, one de novo signature was removed from the inferred set.

Following refinement, we proceeded with the merging step. Initially, the tool detected 14 clusters among the breast cancer samples. We then applied an iterative merging function, resulting in 5 clusters (Additional file 1: Fig. S9), using a cutoff value of 0.8.

Later, we mapped the de novo signatures to known signatures from the COSMIC catalogue and the findings from Degasperi et al.; within the SBS context, 8 de novo signatures corresponded to COSMIC signatures SBS7a, SBS9, SBS17b, SBS26, SBS40a, SBS44, SBS90, and SBS98. Additionally, one de novo signature aligned with the common signature SBS8, and one signature matched the rare signature SBS97, both from the Degasperi et al. study. In the DBS context, we mapped 2 de novo signatures to COSMIC signatures DBS14 and DBS2. Ultimately, one de novo signature, SBSD12 remained unmapped (Additional file 1: Fig. S10), which exhibits a pronounced pattern of C>G and C>T transitions. The detailed signature exposures across the various clusters are presented in (Additional file 1: Fig. S11).

#### Lung cancer analysis

We analysed 1,396 lung tumour samples obtained from the GEL, ICGC, and HMF cohorts. For our study, we selected the following SBS signatures: SBS1, SBS2, SBS3, SBS4, SBS5, SBS8, SBS13, SBS17, SBS18, SBS31, and SBS92, as well as the DBS signatures DBS2, DBS5, DBS13, and DBS20. These signatures collectively formed the reference catalogue for our lung data.

The results from the bNMF analysis revealed 19 SBS signatures (11 of which were identified de novo) and 6 DBS signatures (2 of which were de novo). Following a refinement step, we identified 15 SBS signatures (7 de novo) and maintained 6 DBS signatures (2 de novo). During the refinement process, it was determined that 4 de novo signatures could be attributed to existing signatures (both de novo and input reference catalogue) based on a cosine similarity threshold exceeding 0.9 between the reconstructed and original signatures. After the refinement, we proceeded to the merging phase, where the tool initially identified 15 clusters among the lung cancer samples. We then applied a merging function that iteratively combined these clusters until we arrived at 7 distinct clusters (Additional file 1: Fig. S12), using a cutoff value of 0.8.

Later, we aligned the de novo mutational signatures with known signatures from the COSMIC database and the findings of Degasperi *et al*. In the context of SBS, we identified 4 de novo signatures that corresponded to COSMIC signatures SBS8, SBS17b, SBS44 and SBS92; and 3 de novo signatures that matched signatures SBS107, SBS123 and SBS127 from Degasperi *et al*. For DBS, we successfully mapped two signatures to COSMIC catalogue signatures DBS1 and DBS6. The detailed signature exposures across the various clusters in lung cancer type are presented in (Additional file 1: Fig. S13).

#### Colorectal cancer analysis

Our study examined 2,845 colorectal tumour samples from the GEL, ICGC, and HMF cohorts. We utilised an input reference catalogue comprising 13 SBS signatures SBS1, SBS2, SBS3, SBS5, SBS8, SBS13, SBS17, SBS18, SBS35, SBS88, SBS93, SBS121, SBS157 and 4 DBS signatures DBS2, DBS5, DBS13, DBS20 for the analysis of colorectal tumour data.

Initial bNMF yielded 32 SBS signatures (24 de novo) and 8 DBS signatures (4 de novo). Subsequent refinement reduced these to 24 SBS signatures (16 de novo) and 8 DBS signatures (4 de novo), as 8 de novo signatures were found to be explicable by other signatures, with cosine similarities exceeding 0.9. Post-refinement the algorithm initially identified 15 clusters, which were reduced to 8 distinct clusters through iterative merging with a cutoff value of 0.8 (Additional file 1: Fig. S14).

We compared the de novo mutational signatures identified in our analysis to known signatures from the COSMIC database and the work of Degasperi et al. In the context of SBS, we found 9 de novo signatures aligned with COSMIC signatures SBS10a, SBS10b, SBS15, SBS17b, SBS26, SBS28, SBS44, SBS52 and SBS93. Additionally, one de novo signature matched signature SBS97 from the rare signatures in Degasperi et al. For DBS context, we successfully mapped three signatures to COSMIC signatures DBS3, DBS8, and DBS14, and one de novo to signature DBS25 from the rare category in the Degasperi et al. study.

Ultimately, 6 de novo signatures remained unmapped (Additional file 1: Fig. S15). SBSD12 exhibits a higher density of C>T and T>C mutations, while SBSD13’s density is mostly distributed in C>A, C>T, T>A and T>C. SBSD16 primarily displays density in C>T, T>A, and T>C mutations. SBSD6 has its density distributed across C>A and C>T transitions. SBSD6 shows a high density in C>A and C>T contexts. SBSD8 is the most widely distributed signature among the others, with higher densities in C>A, C>T, and T>A. Finally, SBSD9 has the majority of its density concentrated in C>A, T>C, and T>G transitions. The detailed signature exposures across the various clusters in this cohort are presented in (Additional file 1: Fig. S16).

#### Comparative exposure analysis (BASCULE vs. FitMS)

To investigate the mutational processes underlying breast, lung, and colorectal cancer cohorts, we compared the exposures clustering inferred from BASCULE with the ones estimated with FitMS [23, 27], focusing on single-base substitution (SBS) and double-base substitution (DBS) signatures. Below, we compare the clustering results for each cancer type, highlighting similarities and differences in the identified mutational patterns.

Clustering of breast cancer patients (Additional file 1: Fig. S17) based on exposure profiles from BASCULE and FitMS revealed both shared and distinct patterns, where the median cosine similarity between patients signature exposure is 0.85. The BASCULE-derived cluster G0 and the FitMS-derived cluster G2 exhibited highly similar signature profiles in both SBS and DBS contexts, predominantly driven by APOBEC-related signatures (SBS2, SBS13, and DBS11). This strong association with APOBEC-driven mutational processes suggests a common underlying mechanism in these clusters. In contrast, BASCULE identified a large cluster, G1, characterized by SBS1, SBS90, DBS11, DBS13, and DBS20, indicating a broad mix of mutational processes. FitMS, however, partitioned patients into smaller, more distinct clusters, primarily differentiated by DBS signatures. Notably, FitMS cluster G0 was dominated by DBS20, cluster G4 by DBS13, and cluster G1 by a combination of DBS13 and DBS20.

Analysis of lung cancer patient clustering (Additional file 1: Fig. S18) revealed a prominent group in both BASCULE (G1) and FitMS (G0), strongly associated with tobacco-related mutational processes. These clusters, the largest in each tool’s output, were characterized by elevated SBS4 and DBS2 signatures, consistent with smoking-induced mutagenesis. Although smaller clusters displayed divergent patterns, the cosine similarity in patients exposure profiles between the two tools was 0.94, higher than the similarity in breast and colorectal cancer cohorts. In BASCULE, cluster G0 was distinguished by the presence of DBS1, a signature linked to UV light exposure, which was absent in FitMS results. Additionally, BASCULE cluster G3 and FitMS cluster G6, while differing in patient numbers, shared a common feature: elevated DBS5 signatures associated with chemotherapy-related processes.

For colorectal cancer, clustering (Additional file 1: Fig. S19) with BASCULE and FitMS identified two dominant groups: G1 (BASCULE) and G0 (FitMS). Both groups were characterized by aging-related signatures (SBS1 and SBS5) and DBS20, linked to aristolochic acid exposure, indicating shared underlying mutational processes. A notable distinction was observed in BASCULE cluster G10, marked by the sequencing artifact signature DBS14, which corresponds to SBS28 from the Degasperi study and was prominent in FitMS’s cluster G1. Additionally, BASCULE cluster G3 and FitMS’s cluster G2 shared similar DBS profiles (DBS5 and DBS20) but differed in their SBS signature patterns. Smaller clusters from both tools displayed heterogeneous signature compositions, reflecting the complexity of mutational processes in colorectal cancer, where cosine similarity in patients exposure profiles between two tools was 0.78, indicating the lowest compared to other two tumor types, breast and lung.

#### Skin cancer analysis

Due to the limitation on access to clinical data for the Genomics England, PCAWG and Hartwig Medical Foundation cohorts, our analysis was limited to the ICGC cohort. Analysing 259 samples, two clusters (samples belonging to clusters with less than or equal 20 members eliminated) and 10 mutational signatures (Additional file 1: Fig. S20), include eight established signatures SBS1, SBS3, SBS5, SBS7a, SBS7c, SBS7d, SBS18, SBS23 and SBS43 from COSMIC and Degasperi study, alongside one de novo signatures SBSD4 detected using the BASCULE framework. Initially 5 de novo signatures were extracted, where post-inference mapping (Additional file 1: Fig. S20B), assigned four of the de novo signatures to known predefined signatures (SBS7d, SBS23 and SBS43 from COSMIC and SBS7c from Degasperi study).

Cluster G1, comprising a smaller subset of samples (n=37), exhibits a heterogeneous mutational profile (Additional file 1: Fig. S20C), predominantly enriched in age-related signatures SBS1 and SBS5, along with SBS3 and SBS18 which has been associated with homologous recombination deficiency and damage by reactive oxygen species. This distinct mutational composition suggests an etiology that diverges from the canonical ultraviolet (UV) damage pathway. Additionally, SBS43, classified as a possible artifact in COSMIC, contributes to the mutational landscape of cluster G1, with some samples displaying elevated proportions of this signature. In contrast, cluster G11 (Additional file 1: Fig. S20C), encompassing the majority of samples (n=190), is primarily characterized by the SBS7 family (SBS7a, SBS7c and SBS7d), strongly indicative of UV-induced mutagenesis, aligning with the well-established role of UV exposure in the pathogenesis of skin cancers such as melanoma. Notably, signature SBS23 (from Degasperi catalogue), currently with unknown aetiology, is present, and the contribution of the de novo signature SBSD4 is more pronounced in G11 than in G1. The presence of novel signature SBSD4 identified via BASCULE suggests the existence of previously uncharacterised mutagenic processes within this cohort (Additional file 1: Fig. S20D).

#### Pancreas cancer analysis

We analyzed 343 pancreatic tumor samples using the BASCULE framework, identifying three clusters (samples belonging to clusters with less than or equal 20 members eliminated) and 14 mutational signatures (SBS1, SBS2, SBS3, SBS5, SBS8, SBS13, SBS17b, SBS18, SBS26, SBS31, SBS36, SBS44, SBS101, SBS127) from the COSMIC and Degasperi study catalogues (Additional file 1: Fig. S21A). Seven de novo mutational signatures were initially extracted and subsequently mapped to established signatures through post-inference analysis (Additional file 1: Fig. S21B), including SBS17b, SBS26, SBS36 and SBS44 from COSMIC and SBS8, SBS101, and SBS127 from Degasperi *et al* study.

In cluster G0 (n=278), as the largest subgroup, there is a heterogeneous mutational profile dominated by the age-related SBS1 signature (Additional file 1: Fig. S21C), alongside SBS127 and SBS8 (linked to homologous recombination deficiency, HRD) from the Degasperi catalogue. Cluster G2 (n=39) was characterized by SBS5 (aging-related) and SBS8 (HRD), while cluster G4 (n=26), the smallest subgroup, was defined by SBS1, SBS101, and a complex mix of signatures (Additional file 1: Fig. S21D), rendering it the most intricate subgroup.

#### Esophagus cancer analysis

We examined 315 esophageal tumor samples using the BASCULE framework, identifying two clusters and 14 mutational signatures (SBS1, SBS2, SBS3, SBS5, SBS13, SBS17b, SBS18, SBS28, SBS35, SBS40a, SBS44, SBS122, SBS127, SBSD13) from the COSMIC and Degasperi study catalogues. Kaplan-Meier survival analysis was conducted over an 8-year period, where both clusters indicated nearly identical survival trajectories. This may indicate that the BASCULE framework didn’t stratify samples, with distinguished survival outcomes or that the subgroups are defined by factors unrelated to survival outcomes.

#### Exposure analysis

We utilized the exposure matrix inferred by the tool to analyze mutational signatures associated with clusters. For this purpose we quantified variations in signature exposure distributions among clusters and identified statistically significant enrichment of signatures within specific subgroups compared to other subgroups by generating standardized metrics named Differential Exposure Score [50].

We applied a non-parametric Kruskal-Wallis test and, for each signature, we applied logarithmic inversion for p-values derived from the test. These scores enabled comparative analysis of signature-specific differences across clusters and were visualized using barplots to highlight statistically meaningful distinctions (Additional file 1: Fig. S22). Exposure analysis was conducted across three tumor types, with the results visualized using bar plots and statistical significance assessed at a threshold of p-value=0.05. In the skin tumor cohort (Additional file 1: Fig. S22), signatures such as SBS5 (age-related), SBS7a (UV light-induced), and the de novo signature SBSD8 exhibited significantly different exposures between two distinct clusters. In pancreatic tumors (Additional file 1: Fig. S22), a higher number of clusters revealed a broader set of significantly differentially exposed signatures, including SBS1, SBS2, SBS5, SBS8, SBS31, SBS101, and SBS127. These distinct exposure patterns were reflected in clear patient stratification, as supported by Kaplan–Meier survival analysis. In contrast, analysis of esophageal tumors (Additional file 1: Fig. S22) identified signatures SBS17b, SBS127, SBS18, SBS1, SBSD13, SBS28, SBS40a, and SBS5 as significantly differentially exposed across clusters; however, these differences did not correspond to meaningful patient stratification based on Kaplan-Meier survival curves.

#### Unmapped de novo signatures

The de novo signatures had no mutual signatures with a cosine similarity higher than 0.8 compared to known signatures from the Degasperi *et al* study and the COSMIC catalogue. This threshold value is based on heuristics and may yield different results with varying values. To assess the difference between de novo and known signatures, we subsequently mapped these de novo signatures using a threshold lower than 0.8 (Additional file 1: Fig. S8). The results indicate that some known signatures still exhibit similarities close to 0.8 with specific de novo signatures. This comparison highlights the need for further investigation into selecting an appropriate threshold.

---

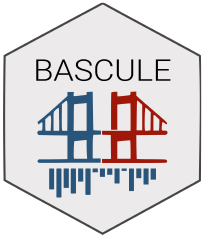

https://caravagnalab.github.io/bascule/

