## Additional file 1 for "BASCULE: Bayesian inference and clustering of mutational signatures leveraging biological priors"

### Additional file 1: Supplementary figures

- Fig. S1: BASCULE performance on synthetic datasets.
- Fig. S2: BASCULE performance comparison on synthetic datasets.
- Fig. S3: BASCULE clustering accuracy on synthetic datasets.
- Fig. S4: BASCULE runtime on synthetic datasets.
- Fig. S5: BASCULE performance comparison on realistic WGS datasets.
- Fig. S6: BASCULE performance comparison on realistic WES datasets.
- Fig. S7: BASCULE clustering accuracy on realistic datasets.
- Fig. S8: Unmapped de novo signatures mapping to Degasperi and COSMIC catalogues in breast and colorectal cancers analysis.
- Fig. S9: Clustering centroids in breast cancer analysis.
- Fig. S10: Unmapped de novo signature in breast cancer analysis.
- Fig. S11: Exposures in breast cancer analysis.
- Fig. S12: Clustering centroids in lung cancer analysis.
- Fig. S13: Exposures in lung cancer analysis.
- Fig. S14: Clustering centroids in colorectal cancer analysis.
- Fig. S15: Unmapped de novo signatures in colorectal cancer analysis.
- Fig. S16: Exposures in colorectal cancer analysis.
- Fig. S17: BASCULE and FitMS exposures in breast cancer analysis.
- Fig. S18: BASCULE and FitMS exposures in lung cancer analysis.
- Fig. S19: BASCULE and FitMS exposures in colorectal cancer analysis.
- Fig. S20: BASCULE analysis of skin cancer data.
- Fig. S21: BASCULE analysis of pancreas cancer data.
- Fig. S22: Differential exposure scores.

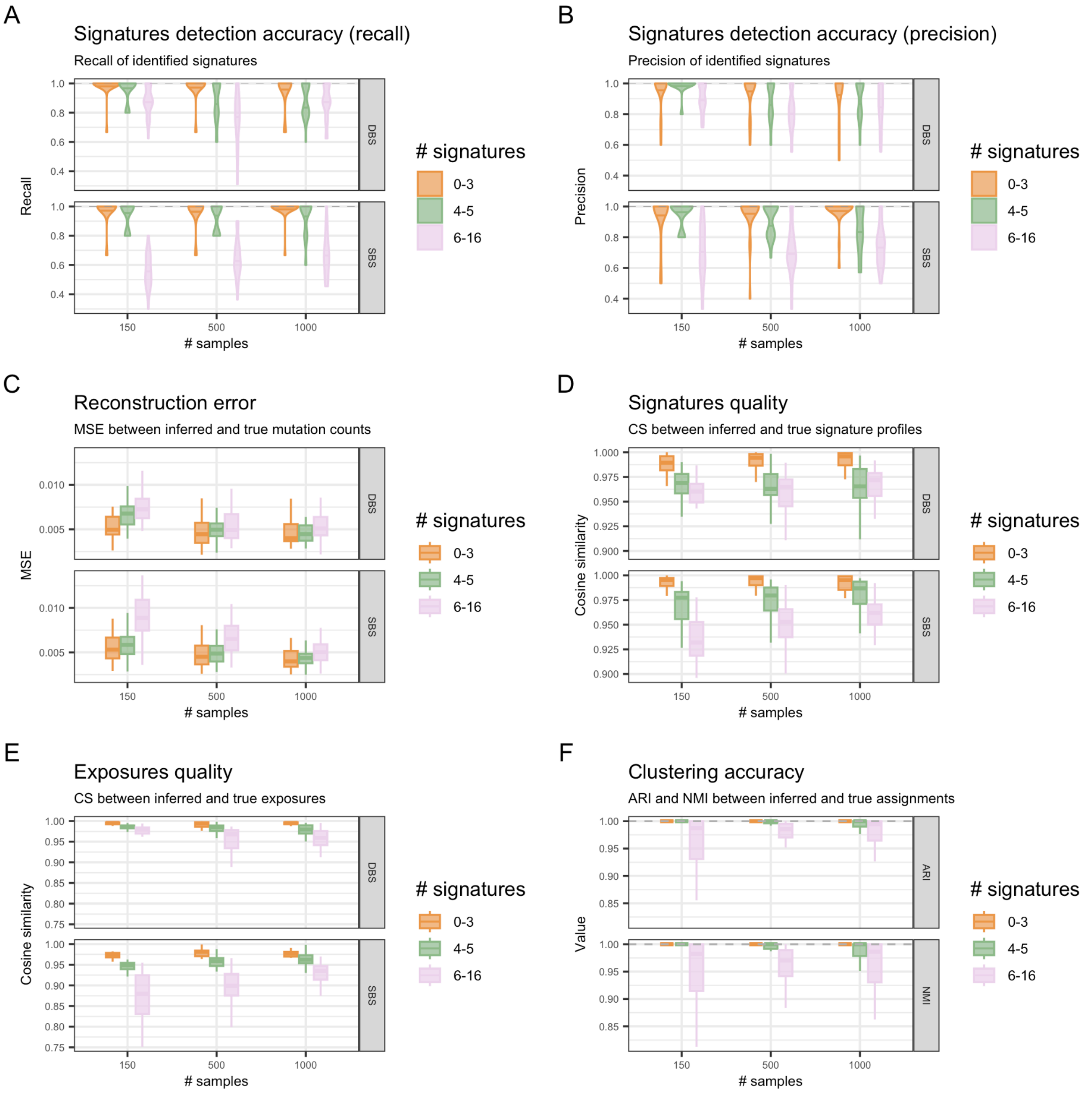


**Fig. S1. BASCULE performance on synthetic datasets.** BASCULE performance assessment with varying the number of signatures in the input dataset. A. Recall computed on the number of retrieved de novo signatures. B. Precision computed on the number of retrieved de novo signatures. C. Reconstruction error (mean squared error) on the input mutation counts matrix. D. Cosine similarity of inferred and true signatures. E. Cosine similarity of inferred and true exposures. F. Adjusted Rand index and normalised mutual information of inferred and true clustering assignments.

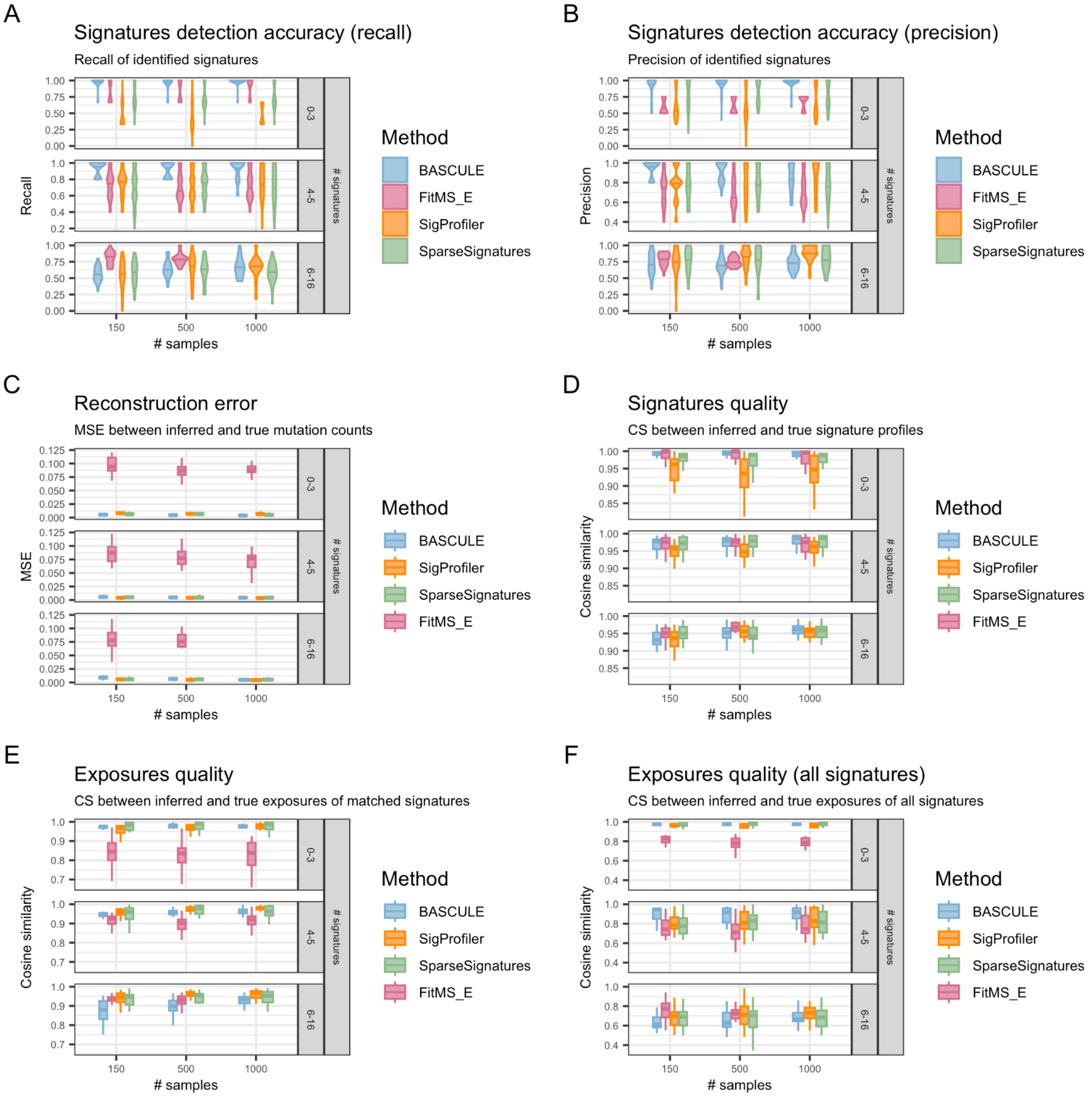


**Fig. S2. BASCULE performance comparison on synthetic datasets.** BASCULE performance compared to SigProfiler, SparseSignatures and FitMS extraction algorithm, with varying the number of signatures in the input dataset. Missing configurations correspond to fits that failed to converge in the specified time. A. Recall computed on the number of retrieved de novo signatures. B. Precision computed on the number of retrieved de novo signatures. C. Reconstruction error (mean squared error) on the input mutation counts matrix. D. Cosine similarity of inferred and true signatures. E. Cosine similarity of inferred and true exposures considering matched signatures. F. Cosine similarity of inferred and true exposures considering matched and unmatched signatures.

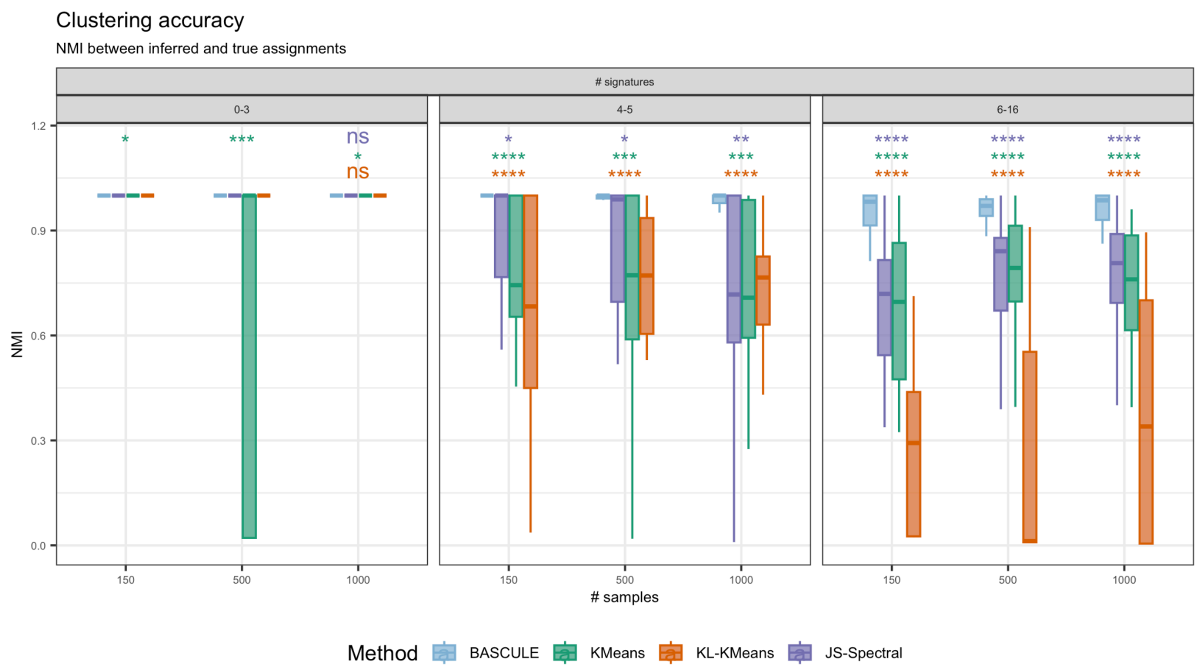


**Fig. S3. BASCULE clustering accuracy on synthetic datasets.** BASCULE clustering performance compared to k-means with Euclidean distance, k-means with Kullback-Leibler distance, and spectral clustering with Jensen-Shannon distance. The p-values reported here are computed to compare BASCULE mean NMI with the competitors mean NMI, using a Wilcoxon test.

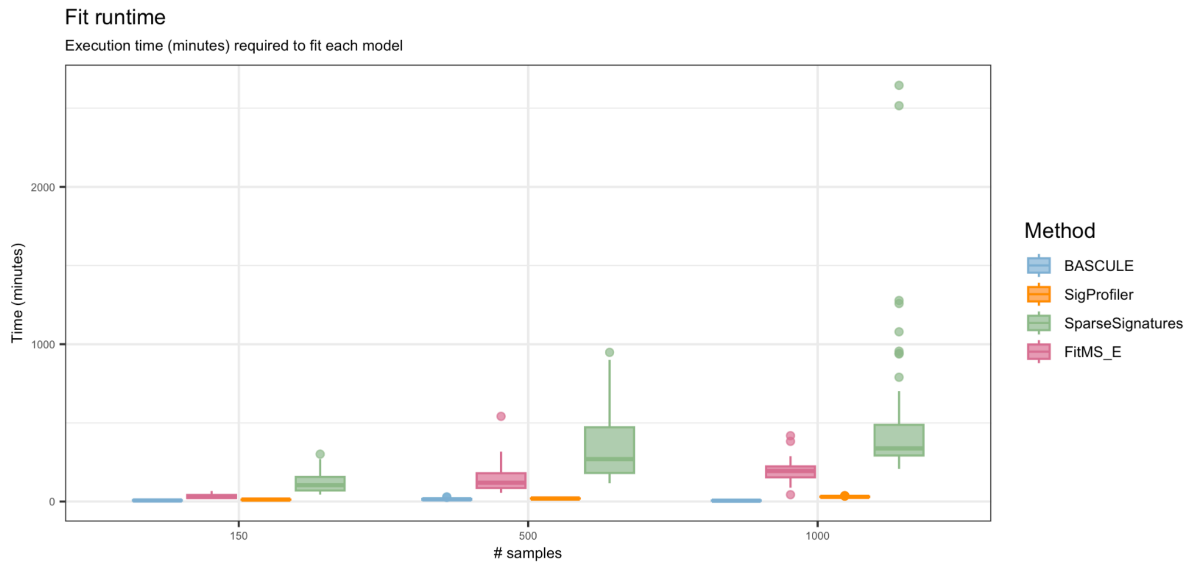


**Fig. S4. BASCULE runtime on synthetic datasets.** Runtime (in minutes) of BASCULE compared to SigProfiler, SparseSignatures and FitMS extraction algorithm.

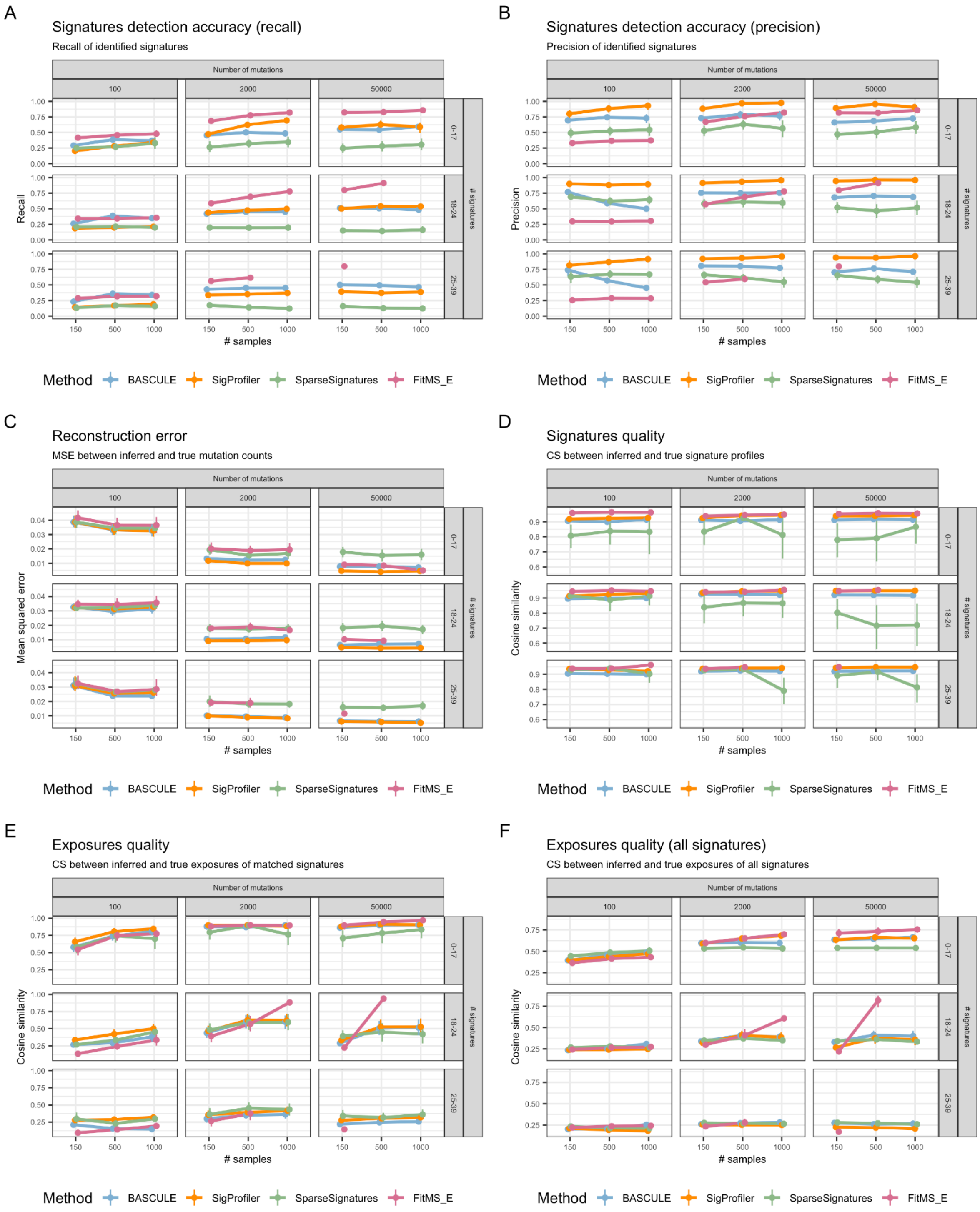


**Fig. S5. BASCULE performance comparison on realistic WGS datasets.** BASCULE and competitors NMF performance on simulated realistic WGS datasets. Missing configurations correspond to fits that failed to converge in the specified time. A. Recall computed on the number of retrieved de novo signatures. B. Precision computed on the number of retrieved de novo signatures. C. Reconstruction error (mean squared error) on the input mutation counts matrix. D. Cosine similarity of inferred and true signatures. E. Cosine similarity of inferred and true exposures considering matched signatures. F. Cosine similarity of inferred and true exposures considering matched and unmatched signatures.

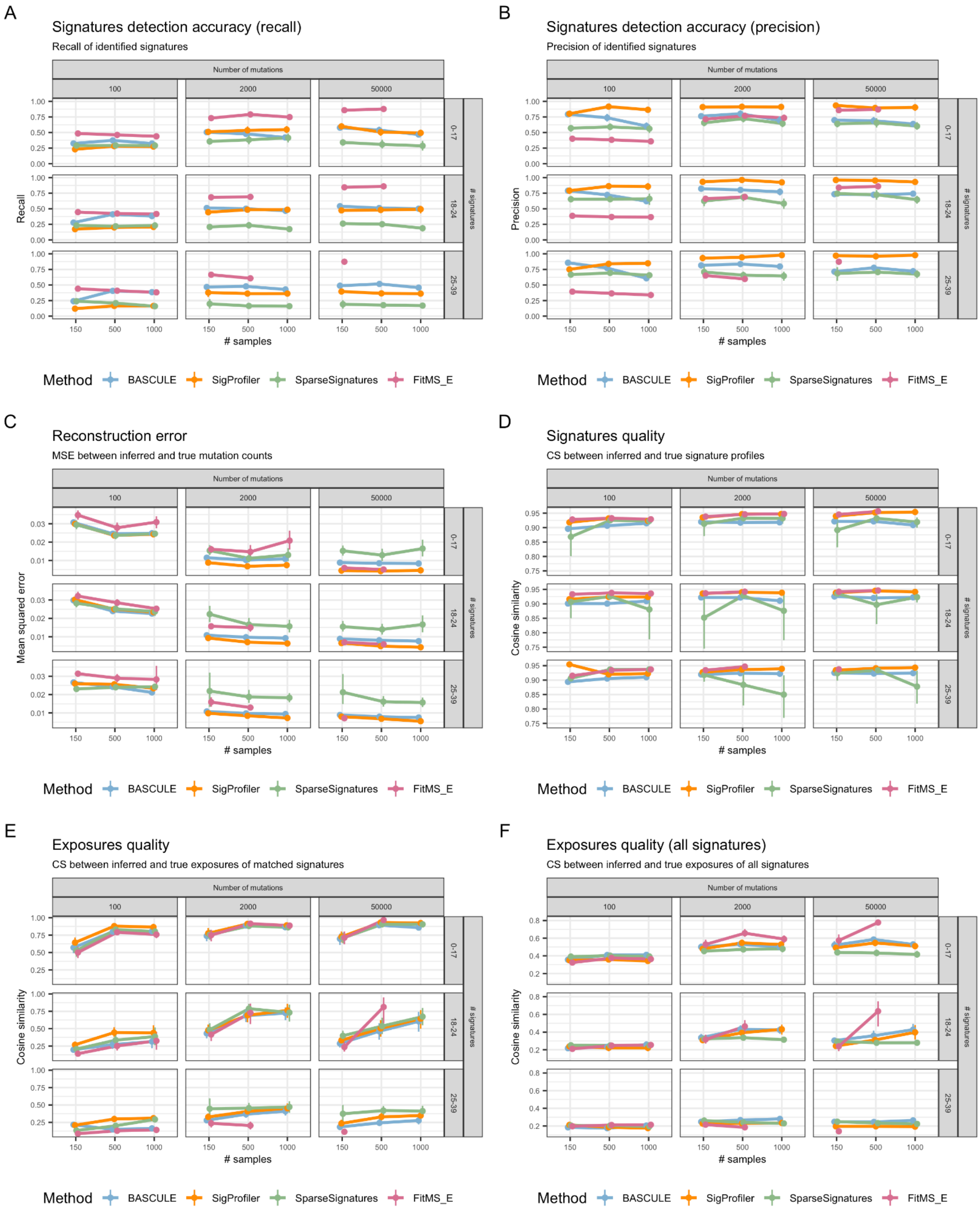


**Fig. S6. BASCULE performance comparison on realistic WES datasets.** BASCULE and competitors NMF performance on simulated realistic WES datasets. Missing configurations correspond to fits that failed to converge in the specified time. A. Recall computed on the number of retrieved de novo signatures. B. Precision computed on the number of retrieved de novo signatures. C. Reconstruction error (mean squared error) on the input mutation counts matrix. D. Cosine similarity of inferred and true signatures. E. Cosine similarity of inferred and true exposures considering matched signatures. F. Cosine similarity of inferred and true exposures considering matched and unmatched signatures.

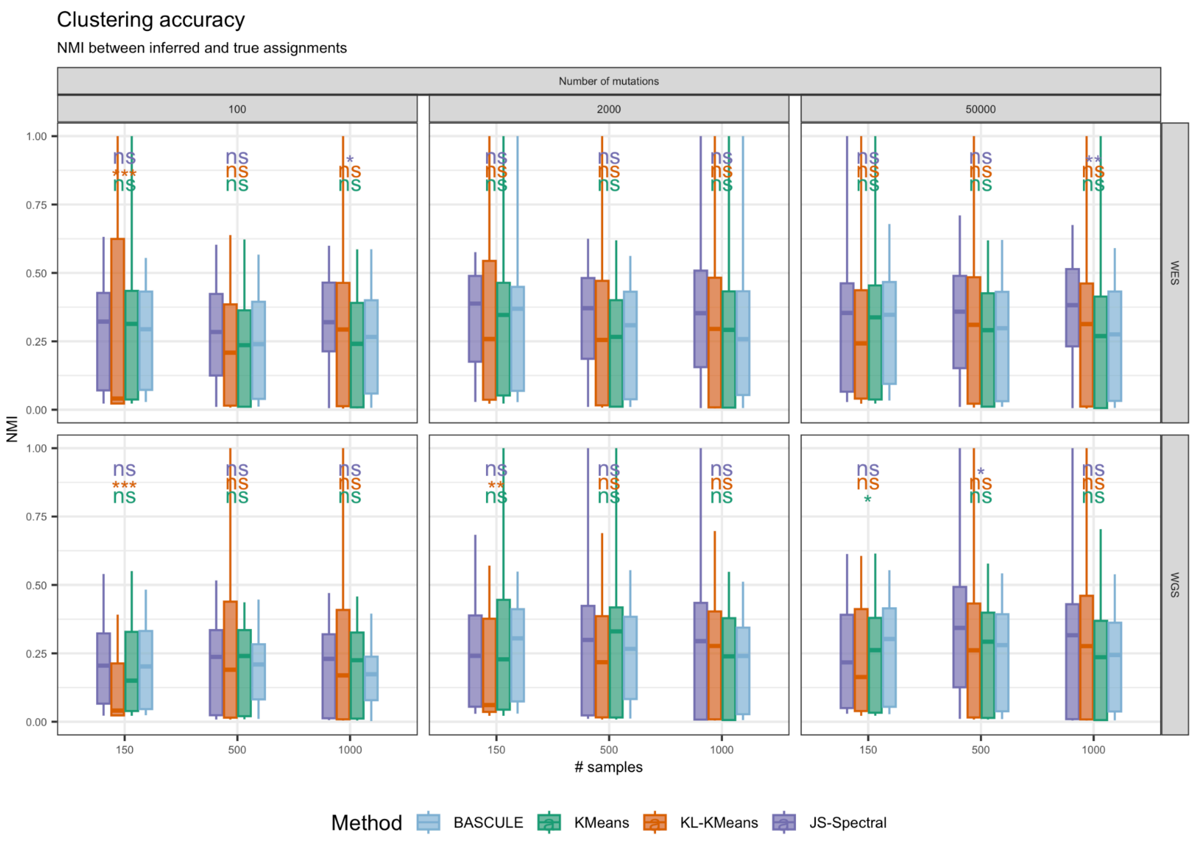


**Fig. S7. BASCULE clustering accuracy on realistic datasets.** BASCULE and competitors clustering performance on the batch of realistic simulated datasets. The plot reports Normalised Mutual Information (NMI) of BASCULE, k-means with Euclidean distance, k-means with Kullback-Leibler divergence (KL-KMeans) and Spectral Clustering with Jensen-Shannon divergence (JS-Spectral). The p-value reported here are computed to compare BASCULE mean NMI with the competitors mean NMI, using a Wilcoxon test.

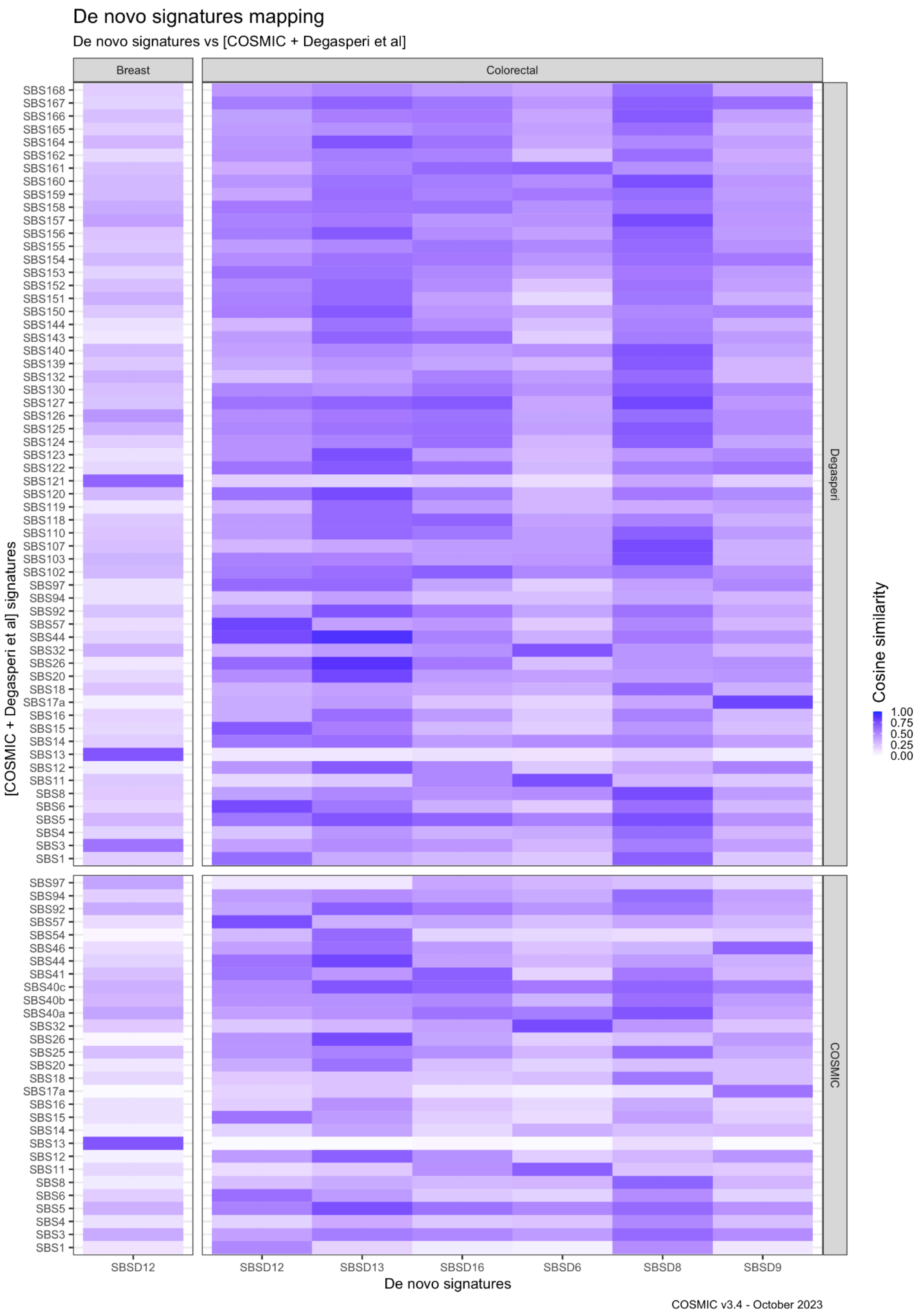


**Fig. S8. Unmapped de novo signatures mapping to Degasperi and COSMIC catalogues in breast and colorectal cancers analysis.** De novo signatures, detected from breast and colorectal tumours, mapped to signatures from the Degasperi *et al.* study and the COSMIC catalogue. Here we report signatures from the reference catalogues with at least one value of cosine similarity greater than 0.6.

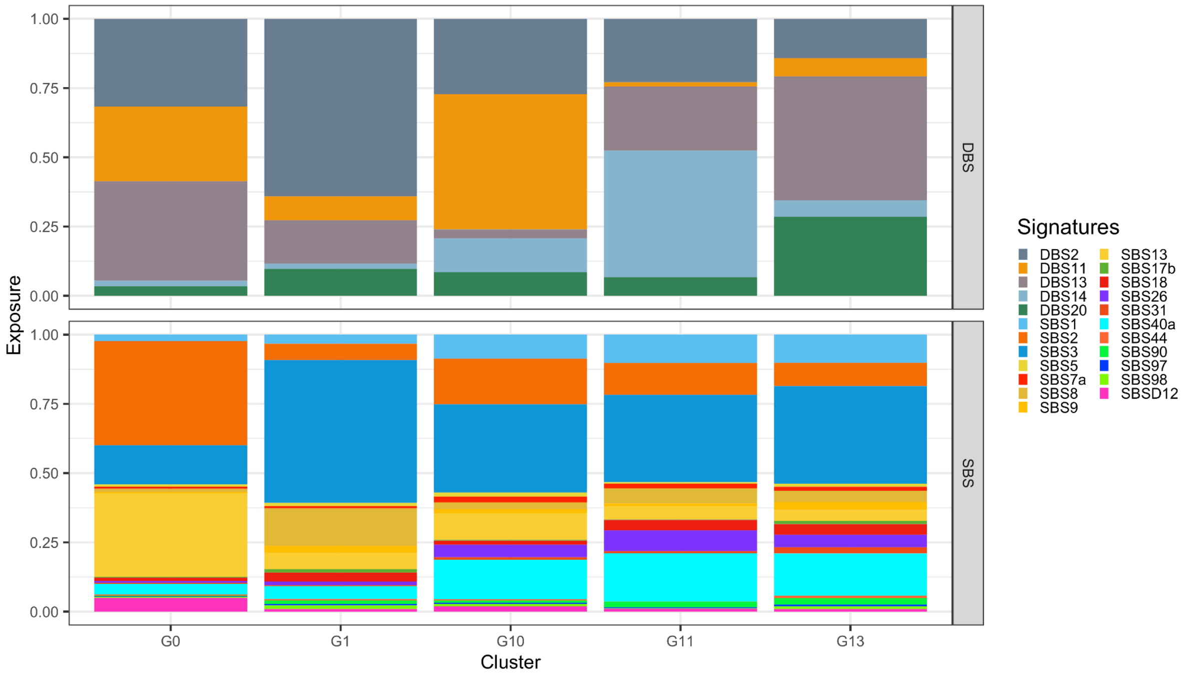


**Fig. S9. Clustering centroids in breast cancer analysis.** Clustering centroids for breast cancers, with 5 clusters and 23 SBS and DBS signatures.

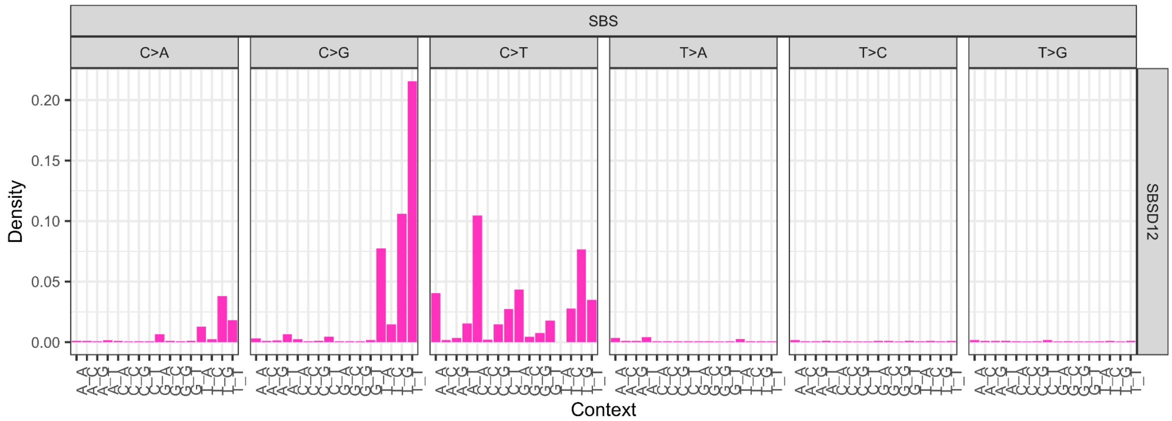


**Fig. S10. Unmapped de novo signature in breast cancer analysis.** Breast cancer de novo signatures that are not mapped to any known signatures from COSMIC or Degasperi et al., based on cosine similarity. See also Fig. S8.

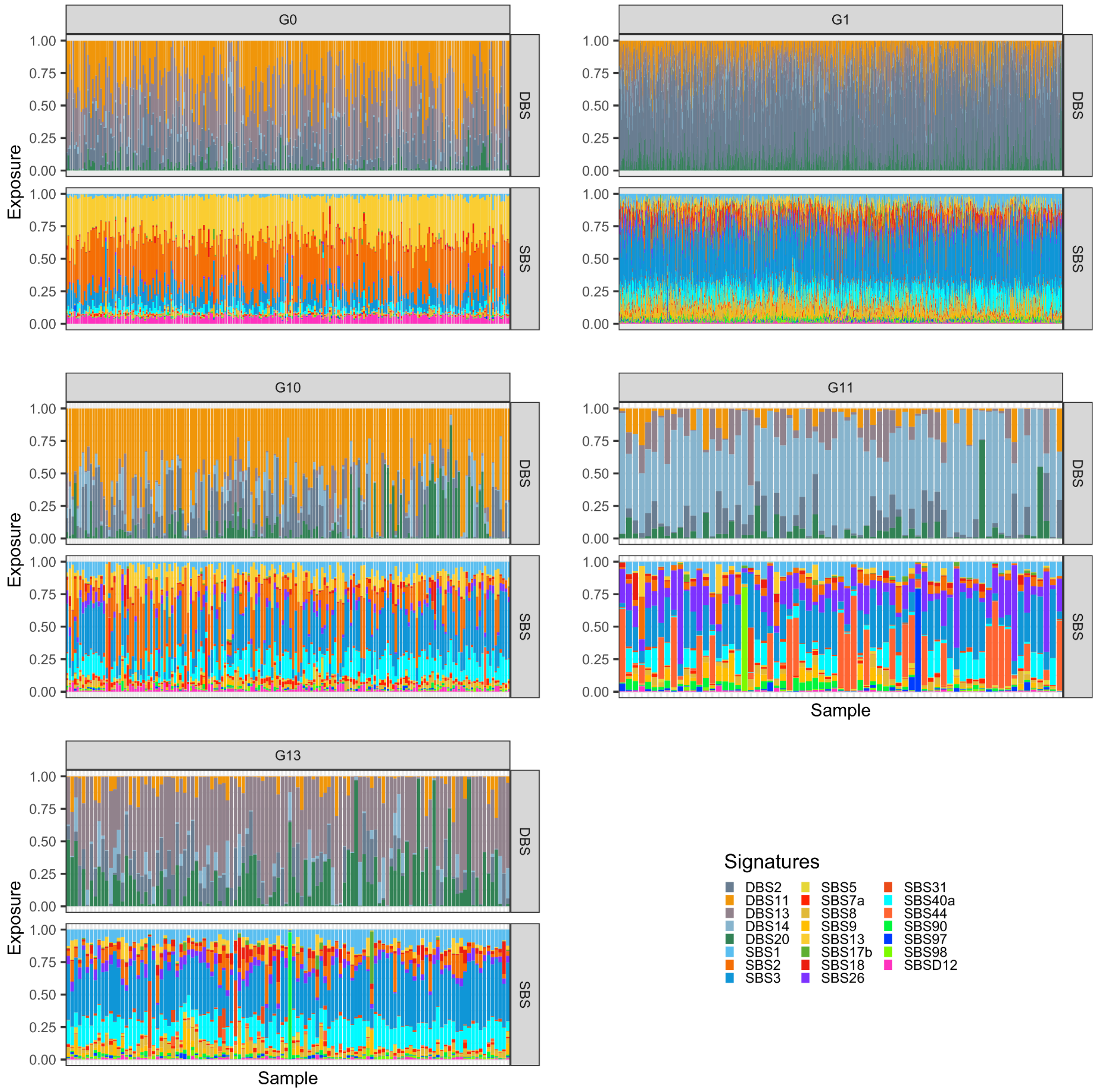


**Fig. S11. Exposures in breast cancer analysis.** The exposure plot of all 5 clusters detected in breast cancers, including both SBS and DBS signatures.

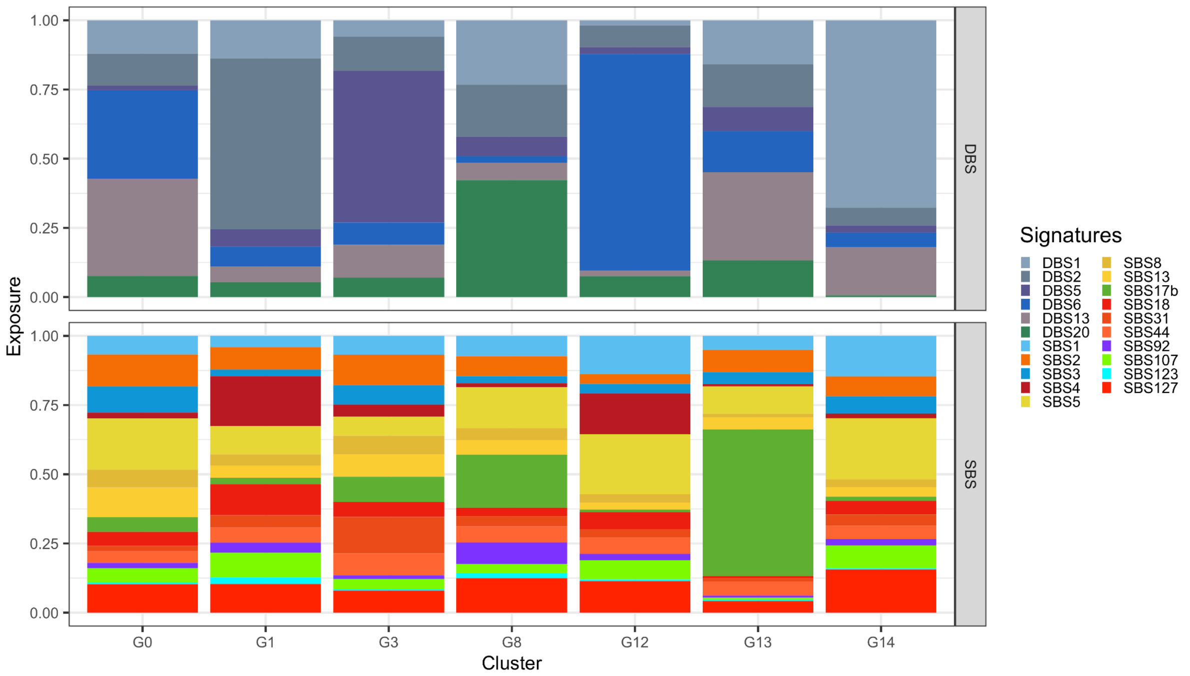


**Fig. S12. Clustering centroids in lung cancer analysis.** Clustering centroids for lung cancers, with 7 clusters and 21 SBS and DBS signatures.

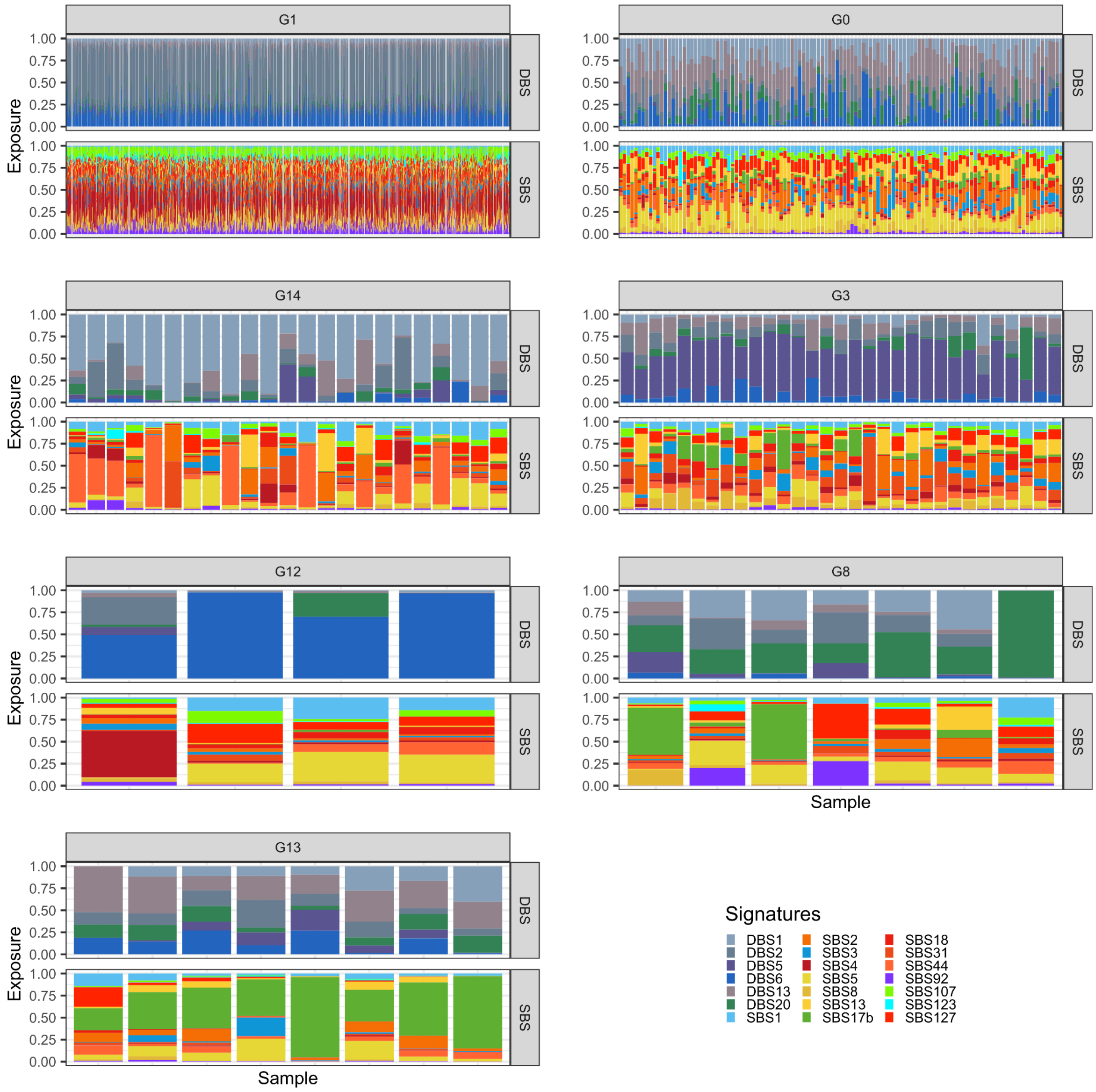


**Fig. S13. Exposures in lung cancer analysis.** The exposure plot of all 7 clusters detected in lung cancers, including both SBS and DBS signatures.

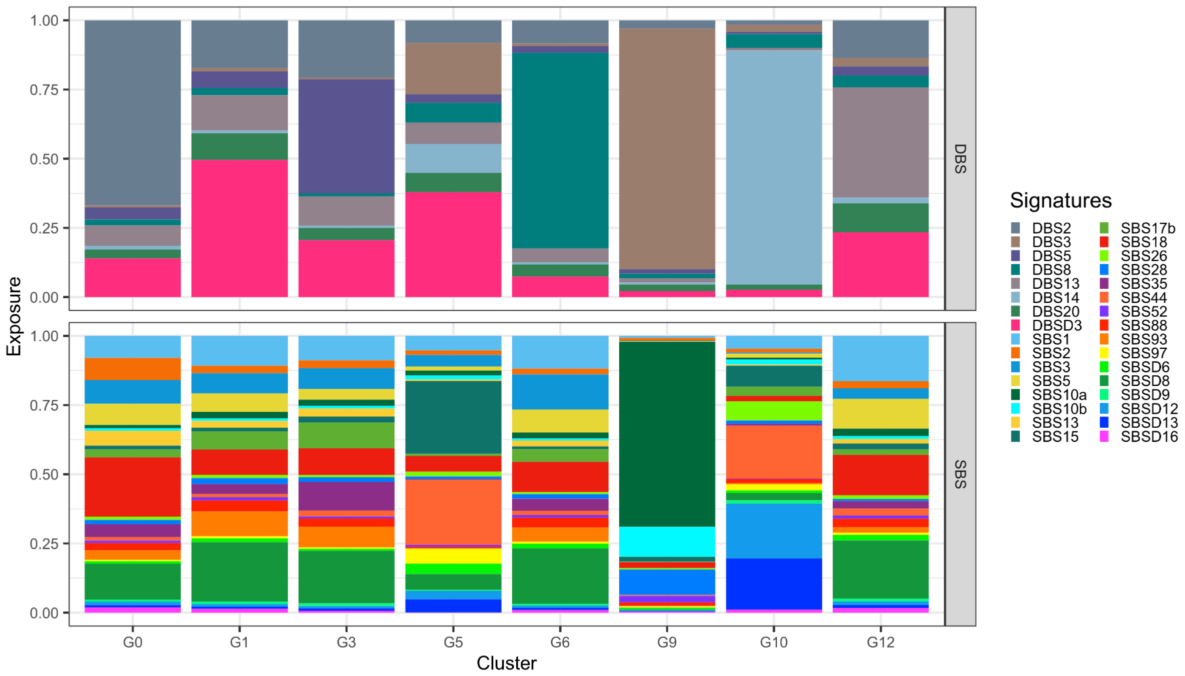


**Fig. S14. Clustering centroids in colorectal cancer analysis.** Clustering centroids for colorectal cancers, with 8 clusters and 32 SBS and DBS signatures.

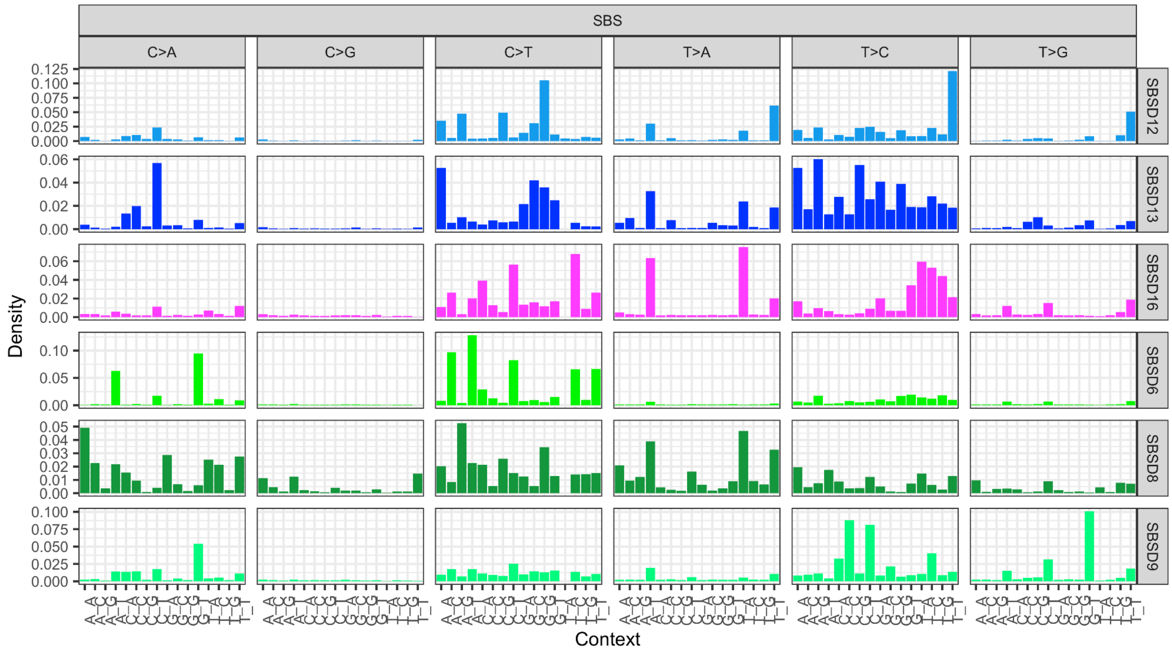


**Fig. S15. Unmapped de novo signatures in colorectal cancer analysis.** Colorectal cancer de novo signatures that are not mapped to any known signatures from COSMIC or Degasperi et al., based on cosine similarity.

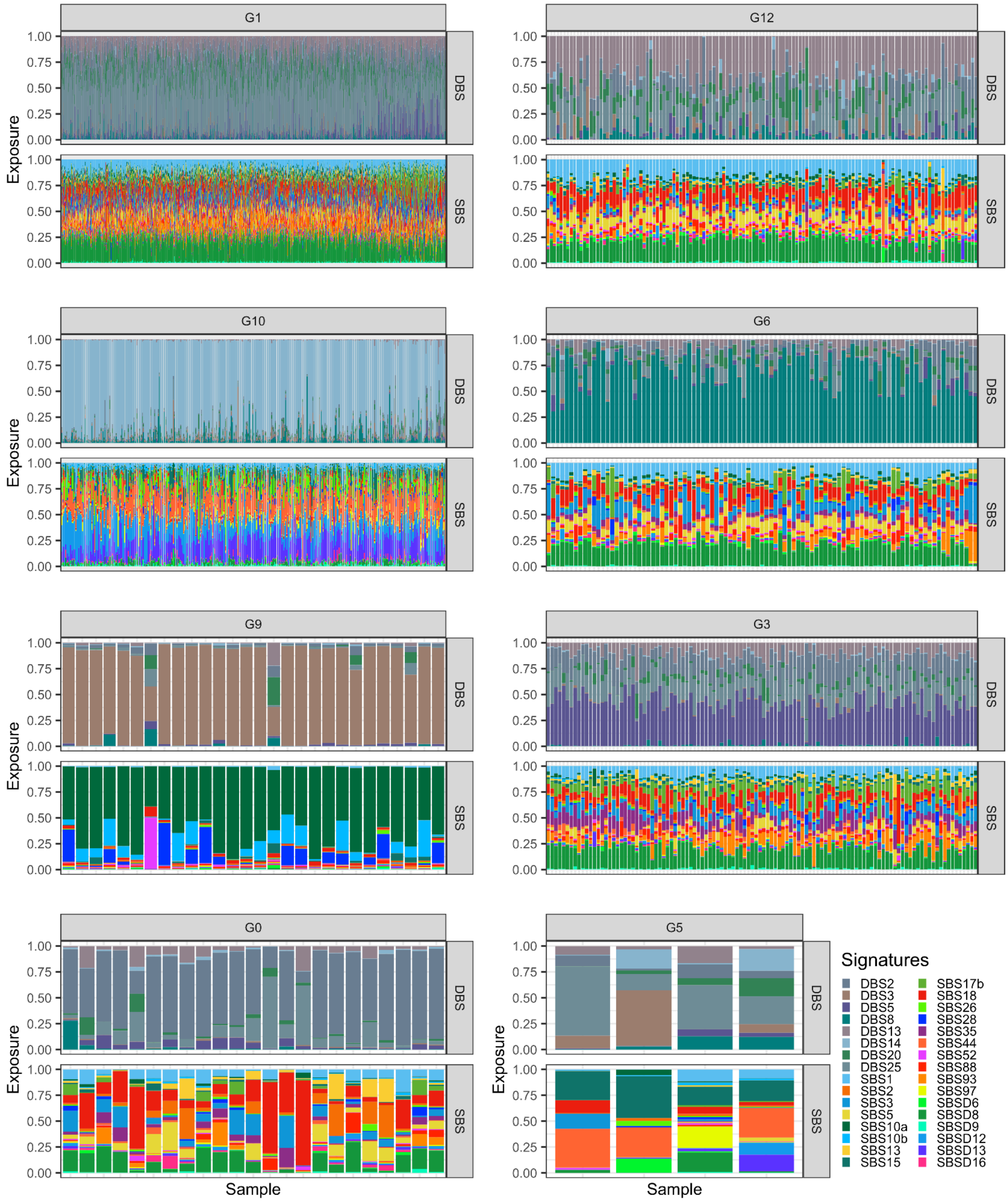


**Fig. S16. Exposures in colorectal cancer analysis.** The exposure plot of all 8 clusters detected in colorectal cancers, including both SBS and DBS signatures.

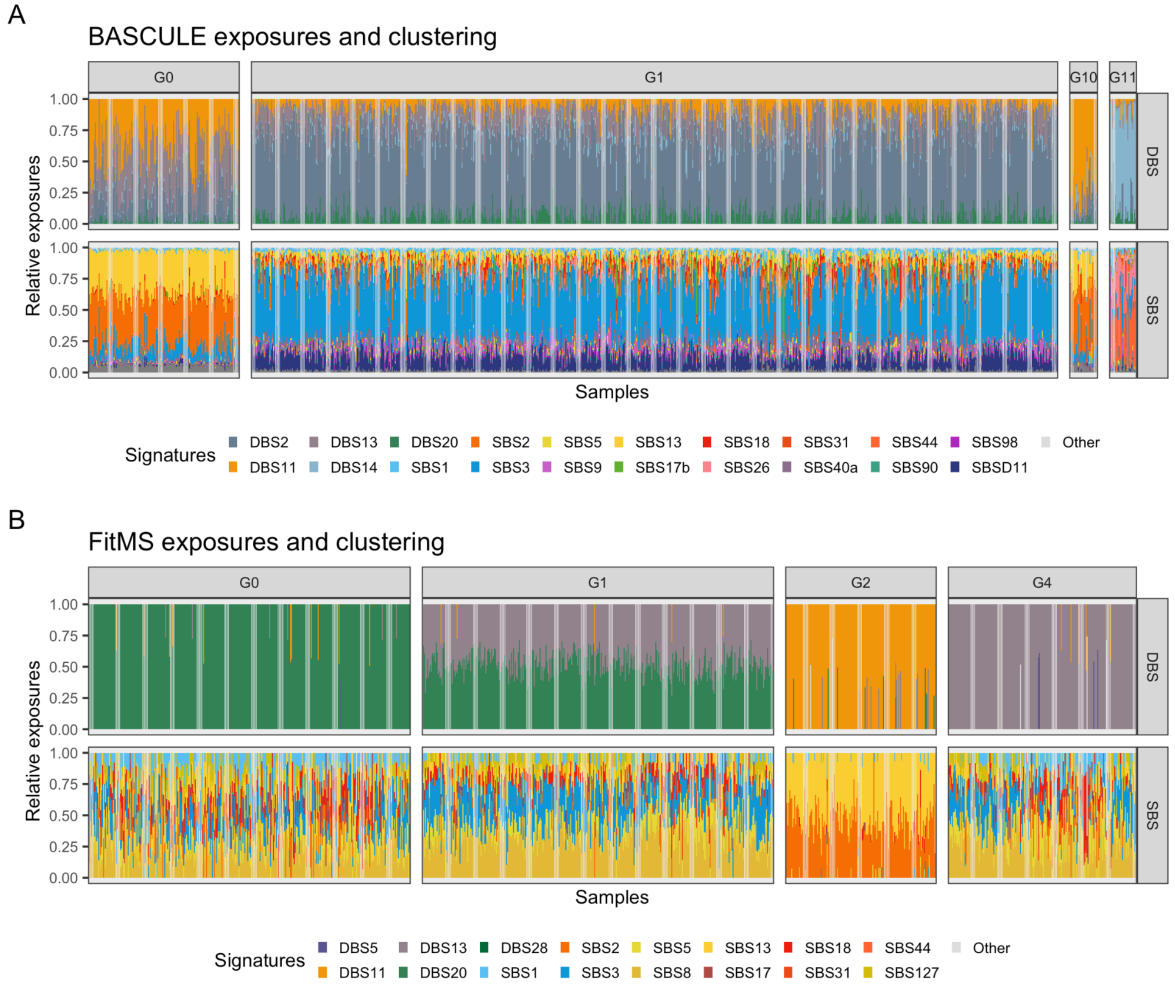


**Fig. S17. BASCULE and FitMS exposures in breast cancer analysis.** BASCULE (A) and FitMS (B) exposures clustered with BASCULE non-parametric clustering model in the breast cancer patients cohort. Only groups with more than 20 patients are shown. Signatures labeled as “Other” have a frequency greater than 0.05 in at least 10 patients.

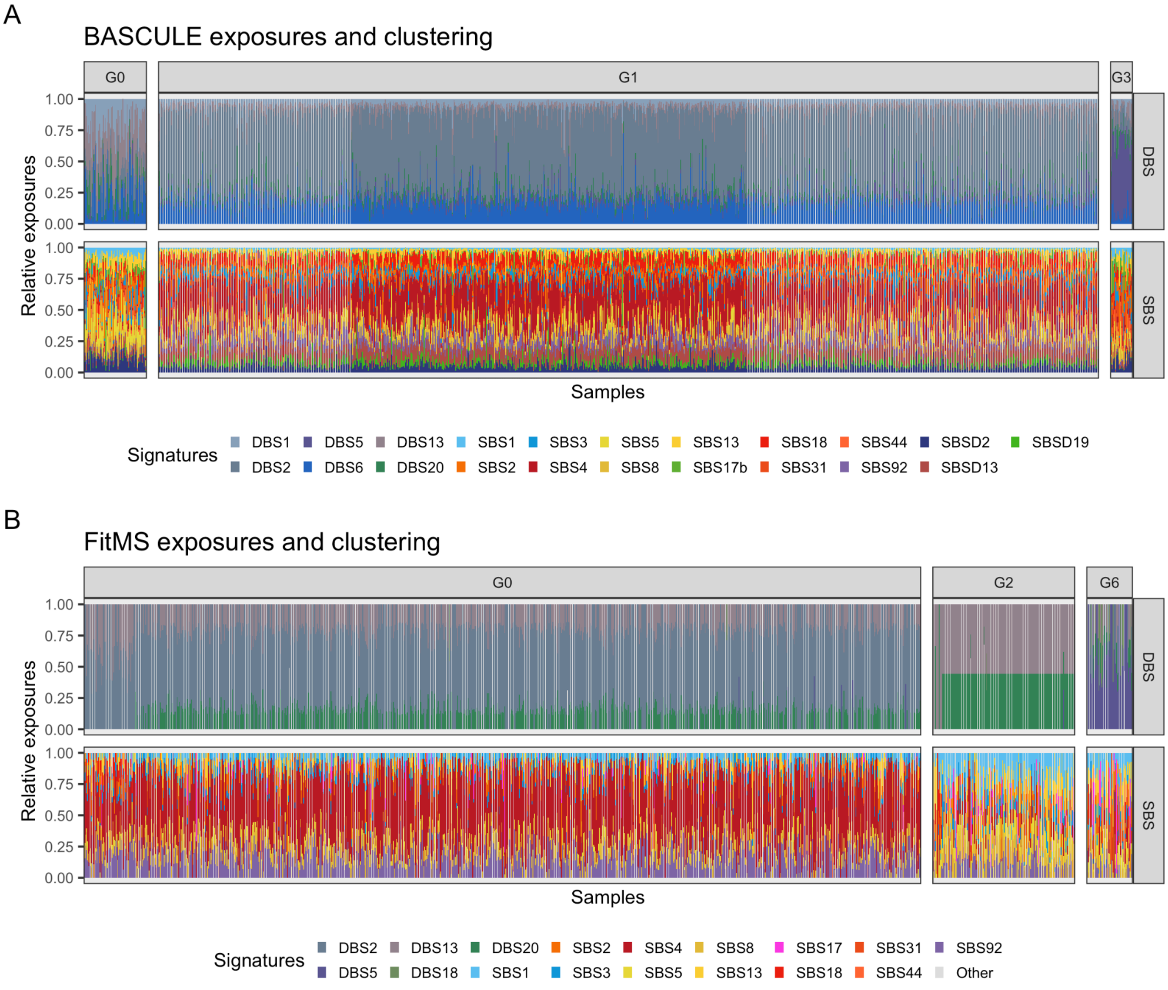


**Fig. S18. BASCULE and FitMS exposures in lung cancer analysis.** BASCULE (A) and FitMS (B) exposures clustered with BASCULE non-parametric clustering model in the lung cancer patients cohort. Only groups with more than 20 patients are shown. Signatures labeled as “Other” have a frequency greater than 0.05 in at least 10 patients.

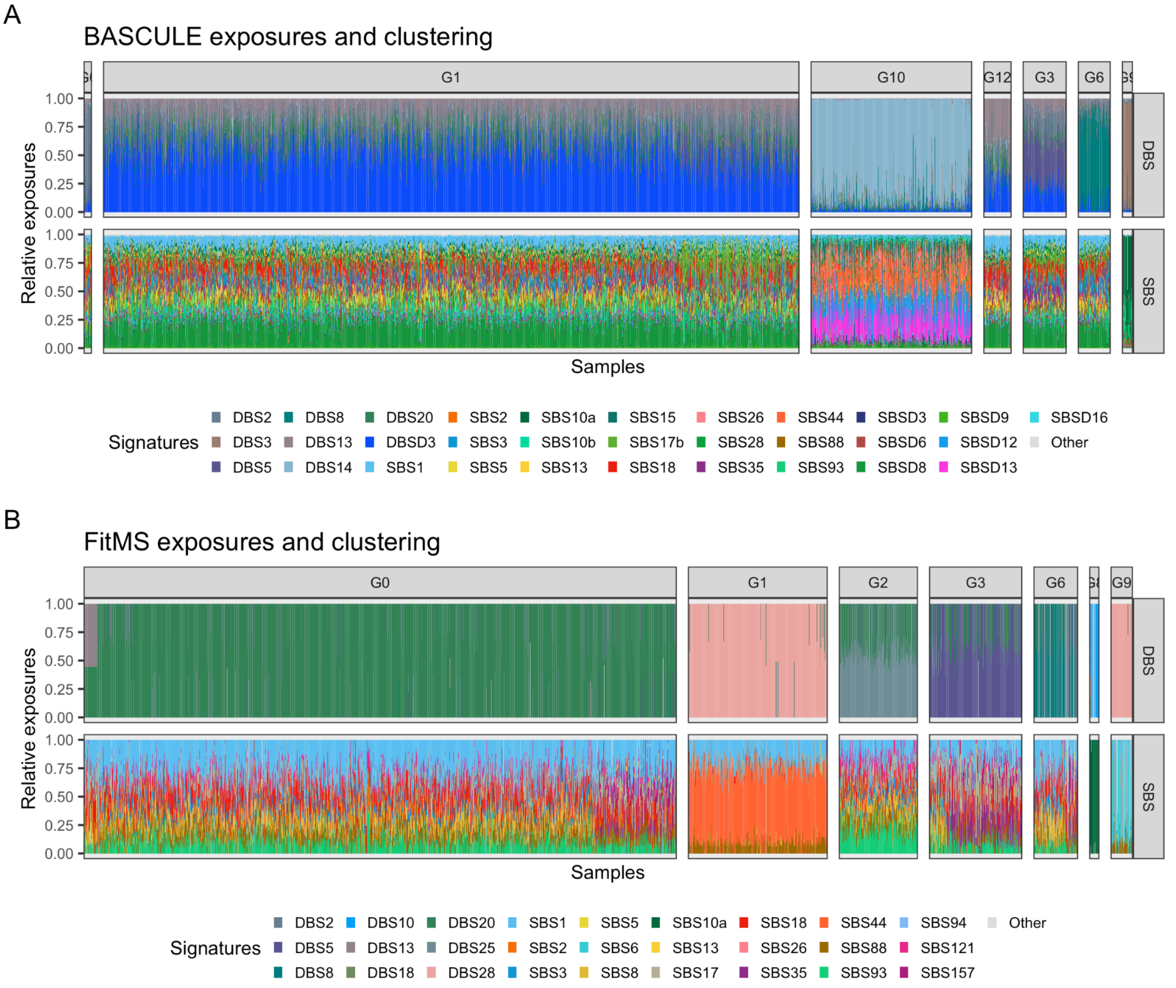


**Fig. S19. BASCULE and FitMS exposures in colorectal cancer analysis.** BASCULE (A) and FitMS (B) exposures clustered with BASCULE non-parametric clustering model in the colorectal cancer patients cohort. Only groups with more than 20 patients are shown. Signatures labeled as “Other” have a frequency greater than 0.05 in at least 10 patients.

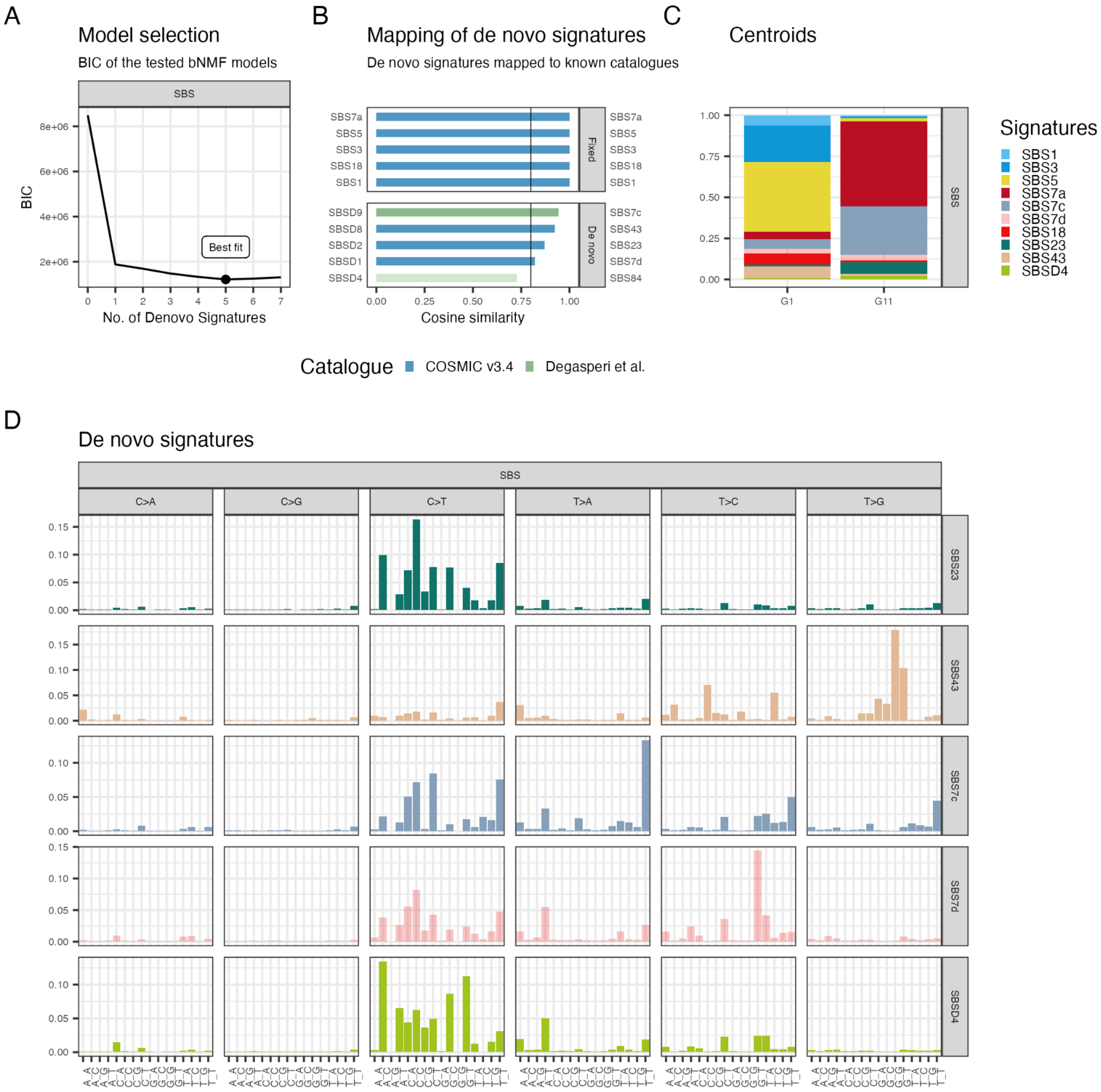


**Fig. S20. BASCULE analysis of skin cancer data.** A. Optimal number of de novo SBS signatures based on BIC criteria, running BASCULE on 259 Skin cancers with a starting catalogue of known signatures, SBS1, SBS3, SBS5, SBS7a, SBS18, SBS99 from COSMIC. B. Mapping for the inferred de novo SBS signatures to known catalogues. SBSD1, SBSD2 and SBSD9 are mapped to SBS7d, SBS137 and SBS7c, respectively. C. Clustering centroids for skin cancers, with 2 clusters and 10 SBS signatures. It includes clusters with more than 20 samples. D. Mutational signatures including the mapped and unmapped de novo signature.

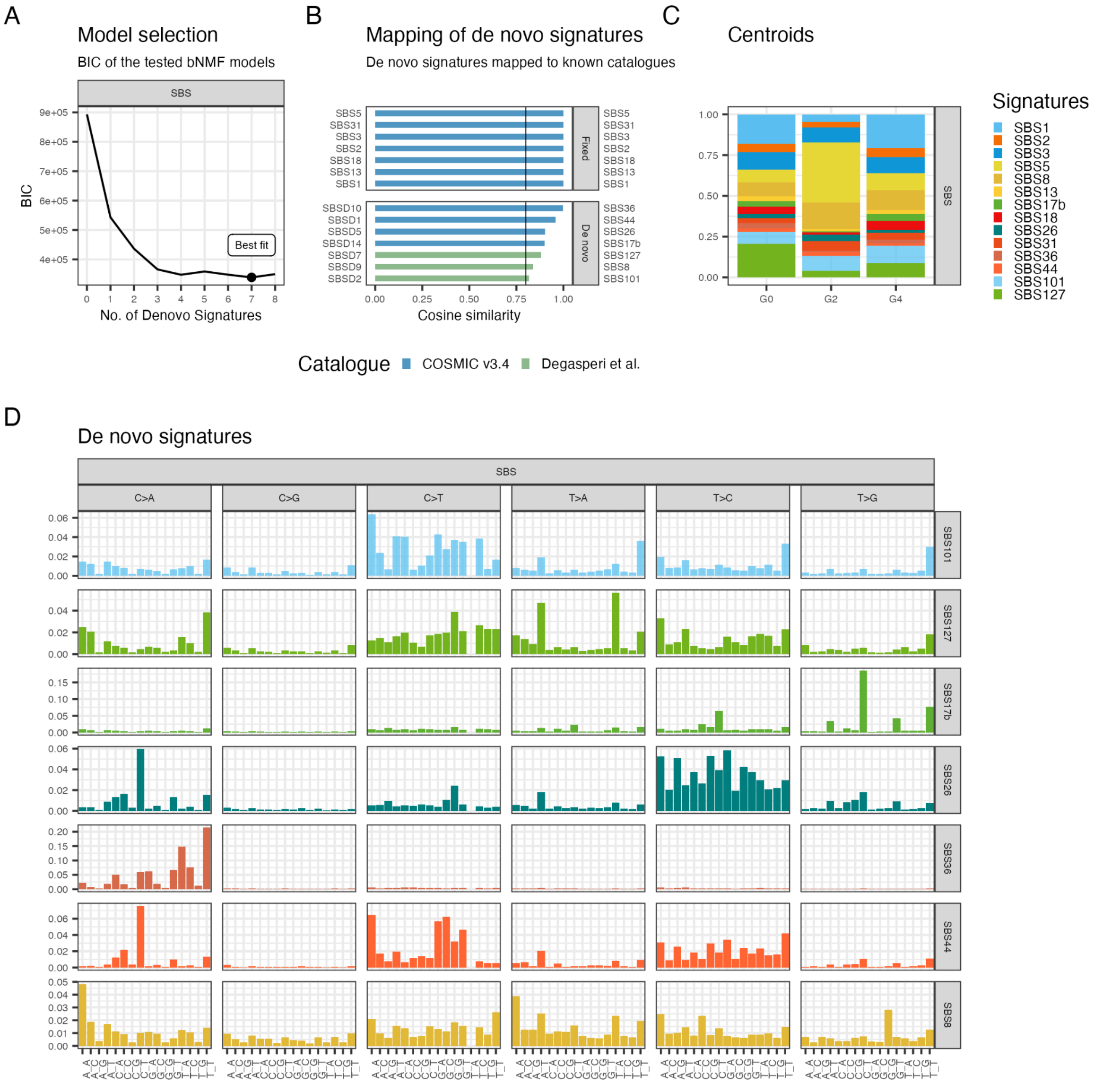


**Fig. S21. BASCULE analysis of pancreas cancer data.** A. Optimal number of de novo SBS signatures based on BIC criteria, running BASCULE on 343 pancreas cancers with a starting catalogue of known signatures, SBS1, SBS2, SBS3, SBS5, SBS13 and SBS18 from COSMIC. B. Mapping for the inferred de novo SBS signatures to known catalogues. SBSD1, SBSD2, SBSD5, SBSD7, SBSD9, SBSD10 and SBSD14 are mapped to SBS44, SBS101, SBS26, SBS127, SBS8, SBS36 and SBS17b, respectively. C. Clustering centroids for pancreas cancers, with 3 clusters and 14 SBS signatures. It includes clusters with more than 20 samples. D. De novo mutational signatures which have been mapped to known signatures from COSMIC and Degasperi study catalogues.

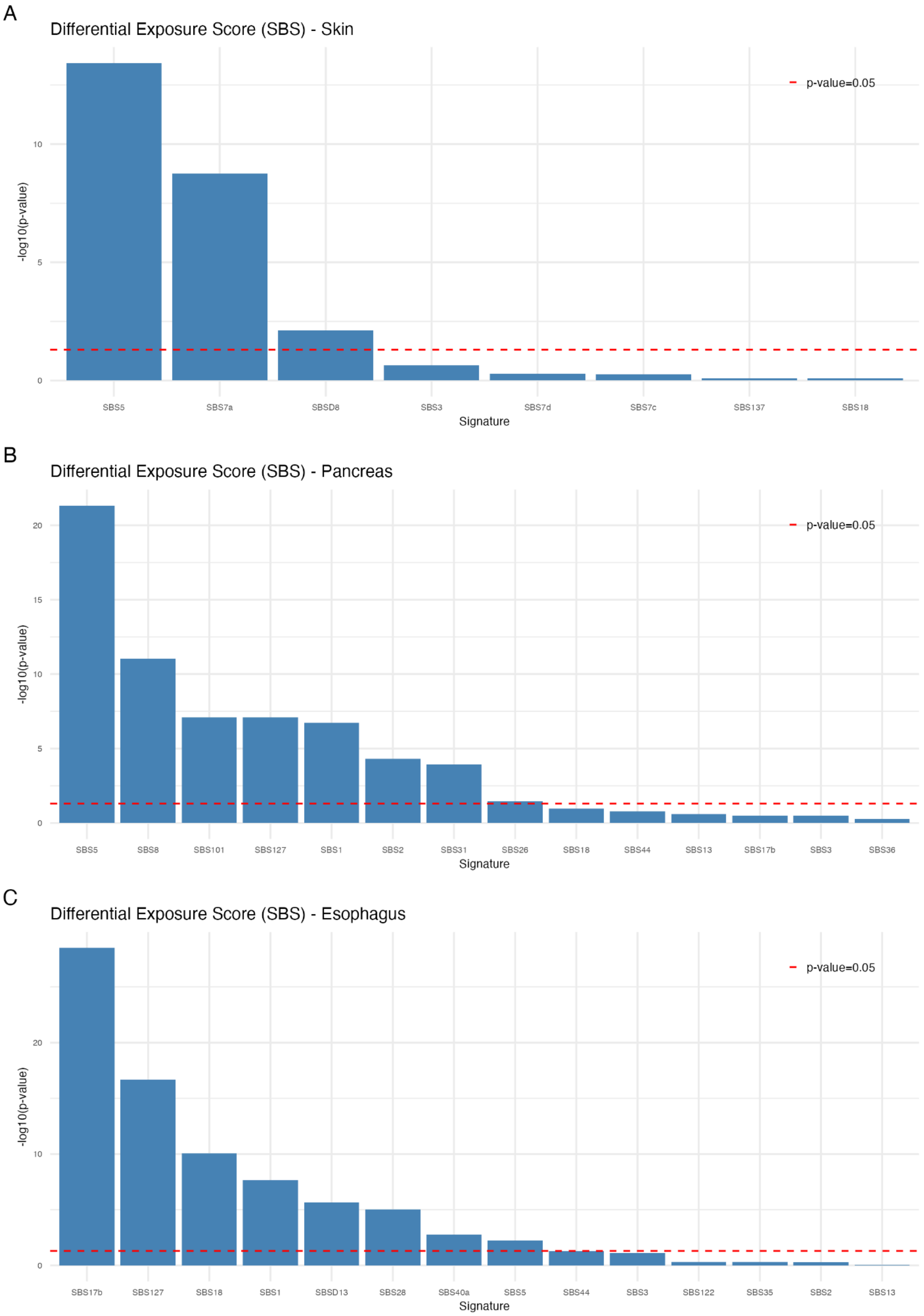


**Fig. S22. Differential exposure scores.** Differential exposure scores plot of three tumor types: skin, pancreatic and esophageal (from top to bottom). The bars are indicating the differential exposure score of the corresponding mutational signature.
